# Subunit-specific conductance of single HCN pacemaker channels at femtosiemens resolution

**DOI:** 10.1101/2024.09.02.610748

**Authors:** Klaus Benndorf, Uta Enke, Debanjan Tewari, Jana Kusch, Haoran Liu, Han Sun, Ralf Schmauder, Christian Sattler

## Abstract

Hyperpolarization-activated cyclic nucleotide-modulated (HCN) channels are tetramers that generate rhythmic electrical activity in neuronal and cardiac pacemaker cells ^1^. The channels are activated by hyperpolarisation of the membrane voltage and additionally tuned by the second messenger cAMP at sympathetic stimulation. There are four mammalian isoforms, HCN1-4 ^2–4^. The single-channel conductance, *γ*, of HCN channels remains debated, with conflicting results ranging from near 1.5 pS for HCN2 ^5,6^ to tens of pS for HCN1 ^7^, HCN2 ^7^ and HCN4 ^7,8^, though the pore structure, viewed to determine the conductance ^9,10^, is either identical or highly conserved. To resolve this controversy, we analyzed all four mouse isoforms mHCN1-4 at femtosiemens resolution. We show that mHCN1, mHCN3 and mHCN4 also generate small conductance values, even smaller than that of mHCN2 with the sequence *γ*_mHCN2_=1.54 pS > *γ*_mHCN1_=0.84 pS > *γ*_mHCN3_=0.54 pS ≍ *γ*_mHCN4_=0.51 pS. As shown by systematic mutagenesis and molecular dynamic simulations, the differences in the conductance are neither generated by the selectivity filter nor the inner gate ^9^, but by defined negative charges in the outer channel vestibule increasing cation occupation. In line with these results, heteromers of mHCN2 with either mHCN1, mHCN3 or mHCN4 lead to graded single-channel currents in-between those of the respective homomeric channels. Our approach at femtosiemens resolution provides insight into the function of recombinant and native HCN channels at the level of single subunits and is thus promising for the development of subunit-specific drugs acting on these clinically highly relevant channels ^11^.

---

Hyperpolarization-activated cyclic nucleotide-modulated (HCN) ion channels evoke the so-called ‘funny’ current *I*_h_ (*I*_f_, *I*_q_) ^4,12^ that generates electrical rhythmicity in many types of neurons ^1^ and pacemaker cardiomyocytes ^13^. In neurons, these channels also contribute to the constraint of long-term potentiation, modulation of the working memory, synaptic transmission as well as resonance and oscillations ^1^.

The primary stimulus for the activation of HCN channels is hyperpolarization of the membrane voltage. The typically slow and sigmoid time course of activation of HCN channels opposes the repolarizing phase of the action potential and generates the slow depolarizing pacemaker potential ^14^. At sympathetic stimulation, the second messenger cAMP binds to the HCN channels, accelerating the electrical rhythmicity by boostering channel activation.

In mammals, four subunit isoforms have been identified, HCN1-4 ^2–4^. All four subunit isoforms can form functional homotetrameric channels ^15–17^. Each subunit contains a cyclic-nucleotide binding domain (CNBD) in the C-terminus for the binding of cAMP ^18^, resulting in four CNBDs per channel. HCN2 (Ludwig et al., 1998) and HCN4 channels (Ludwig et al., 1999) are strongly stimulated by cAMP whereas HCN1 channels are reported to be less sensitive to cAMP ^19^. In contrast, for HCN3 channels no stimulating cAMP effect was found ^17,20^. At heterologous expression, also functional heterotetrameric channels have been identified for HCN1/HCN2 ^21,22^, HCN1/HCN4 ^23^, and HCN2/HCN4 ^24^. Heteromerization was confirmed for all combinations of isoforms by co-immunoprecipitation apart from HCN2/HCN3 ^25^.

Regarding the channel architecture, that of homotetrameric HCN1, HCN3 and HCN4 channels has been elucidated over the last years by cryo-electron microscopy (cryo-EM), both in the presence and absence of bound cAMP ^9,10,26–28^.

The dual activation of HCN channels by voltage and cAMP has been widely studied in ensemble currents from HCN1, HCN2 and HCN4 channels, either in whole cells or macropatches containing typically hundreds of channels. Single-channel currents were identified in sino-atrial pacemaker cells by DiFrancesco decades ago ^12,29^, now known to be preferentially formed by the cardiac isoform HCN4^30^. Compared to the vast majority of ion channels, the conductance is exceptionally small in the range of 1 pS. This makes their analysis technically very challenging. Among recombinant channels, only HCN2 channels have been analyzed at the single-channel level. Their conductance was determined to be ∼1.5 pS ^6^ and 1.67 pS ^5^, confirming the low values for sino-atrial cells. Evidence for a respective low conductance of 0.68 pS for HCN channels was also provided by nonstationary noise analysis in pyramidal neurons ^31^.

In contrast to these studies, larger conductance values by at least an order of magnitude were reported for HCN1, HCN2 and HCN4 channels ^32^, HCN4 channels ^8^ and natural HCN channels in rat hippocampal neurons ^33^, however, without demonstrating the typical slow activation time course. It is very unlikely that both the high and the low conductance values are true ^34^.

Given the controversy over the reported large and small conductance of HCN channels and the prominent roles of these HCN channels in heart and brain function, we performed systematic single-channel analyses of all four HCN isoforms of the mouse (mHCN) in patches of *Xenopus* oocytes expressing these channels heterologously. We demonstrate on the one hand that the single-channel conductance of all homomeric channels is small and aligns with the simulated conductance of mHCN2 channels based on atomistic molecular dynamic (MD) simulations. On the other hand, we found that the single-channel conductance differs significantly among the four isoforms, despite the amino acid sequence in the two narrow pore regions, the selectivity filter and the inner gate ^9^, is either identical or very similar across all isoforms, respectively. By mutagenesis and MD simulations, we identified the atomistic mechanism behind the differing unitary conductance, which is due to different amounts of negative charges in the outer vestibule of the channel pore, resulting in significant differences in the cation occupancy at the extracellular side. Lastly, we demonstrate at the single-channel level that mHCN2 subunits can form functional heteromers with each of the other three mHCN isoforms.

## Results

### Single-channel currents of the four HCN isoforms

In whole oocytes we observed with the TEVC technique that all four isoforms mHCN1-4 express robust currents (Extended Data Fig. 1a) and that both the voltage of half maximum steady-state activation and the slowness of activation follow the sequence mHCN3>mHCN4>mHCN2>mHCN1 (Extended Data Fig. 1b-d). Together with the observed deactivation kinetics (Extended Data Fig. 1e) these data were used to set the conditions for single-channel recordings (Fig. 1) (see Materials and Methods and Supplementary Results).

**Fig. 1.**
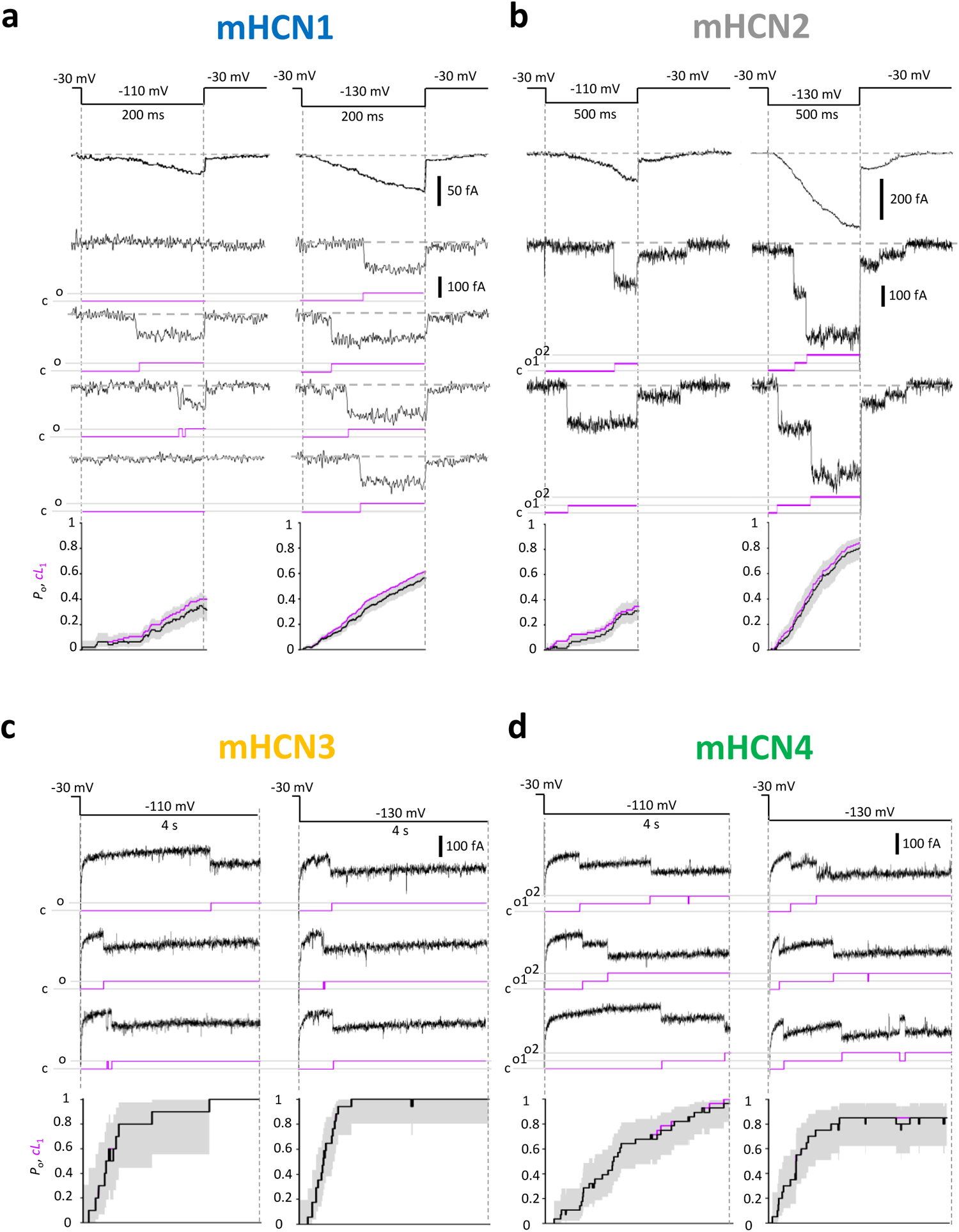
Single-channel currents of wt mHCN channels. Representative traces and patches of the four indicated mHCN isoforms containing either one or two channels. The voltage was -110 mV and - 130 mV. Below each trace, the corresponding idealized traces are shown in magenta; c: closed level; o or o1, o2: open level(s). Bottom: Superimposition of the cumulative first latency *cL*_1_(*t*) (magenta) with the open probability, *P*_o_(*t*) (black). The 95% confidence intervals along *P*_o_(*t*) are indicated in shades of gray. For all isoforms *cL*_1_(*t*) approximately matches *P*_o_(*t*). **a,b**, Fast activating mHCN1 and mHCN2 channels. Filter 200 Hz. The single-channel current traces were corrected for linear leakage and capacitive currents by subtracting nulls (mHCN1: -110 mV: 95 traces, 44 nulls; -130 mV: 371 traces, 108 nulls; mHCN2: -110 mV: 36 traces, 12 nulls; -130 mV: 58 traces, 1 null). The ensemble-average currents (top) as well as the superimposed time courses of the idealized traces (*P*_o_(*t*)) and cumulative first latency, *cL*_1_(t), (bottom) were obtained from these corrected traces. **c,d**, More slowly activating mHCN3 and mHCN4 channels. Filter 100 Hz. The traces were not corrected for linear leakage and capacitive components. For mHCN3, *P*_o_(t) and *cL*_1_(t) were formed from 10 (-110 mV) and 17 (-130 mV) traces whereas for mHCN4 14 (-110 mV) and 10 (-130 mV) traces were included. In case of mHCN3, *P*_o_(t) and *cL*_1_(t) superimpose nearly completely.

As reference we recorded first unitary currents, *i*, from mHCN2 channels (Fig. 1b). To obtain a reasonable number of traces with no openings (nulls) for subtraction of leakage and capacitive currents, short pulses with respect to the full activation time were applied. Both the ensemble average currents (top) and the idealized traces, reflecting the time course of the increasing open probability, *P*_o_(*t*), (bottom) demonstrate the characteristic slow sigmodal activation. The amplitude of the unitary currents was locally measured from time intervals >50 ms before and after an opening transition to minimize effects of possible subtle changes of the leak. The current amplitudes were determined by amplitude histograms (Extended Data Fig. 2a,b). Even the unitary tail currents at 50 fA at -30 mV were reliably resolved (Extended Data Fig. 2c). Using equation (1), a unitary conductance of *γ*_mHCN2_=1.54±0.02 pS was obtained (Table S1). The scatter of the unitary conductance, illustrated by the box plot in Fig. 2a and the line in Fig. 2b, was computed by this value. Superimposition of the cumulative latency until the first opening, *cL*_1_(t), with *P*_o_(*t*) (Fig. 1b, bottom) reveals that *P*_o_(*t*) is mainly a function of *cL*_1_(*t*), indicative that the channels regularly remain open after they have once opened. These results not only confirm previous conductance values in the low range ^5,6^ but also demonstrate that our recording conditions were well suited to reliably study unitary currents down to the range of 50 fA.

**Fig. 2.**
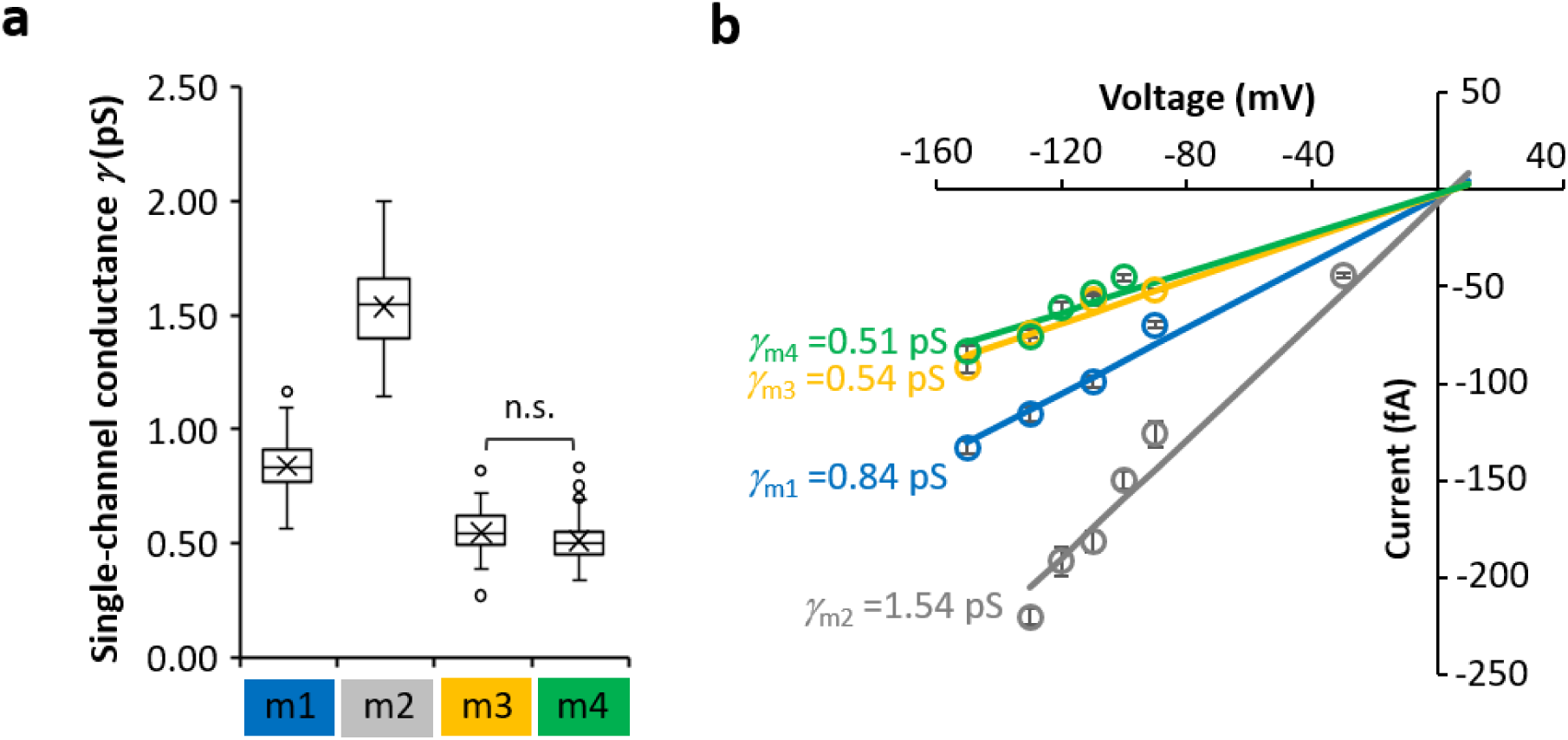
Single-channel conductance of mHCN isoforms. **a**, Boxplot of the unitary conductances. Outliers are included. The plot was generated from individual conductance values obtained by equation (1), taking into account the reversal potential for the used K^+^ concentrations (see Materials and Methods). With P=0.01 all conductance values are significantly different apart those for mHCN3 and mHCN4 (n.s.; pairwise t-tests with Bonferroni-Holm correction). The numbers of included openings, *n*_op_, and patches, *n*_p_, are provided by Table S1. **b**, *iV*-relationships for unitary mHCN currents. The mean conductance values in a were used for calculating the straight lines. The conductance of the channels groups in the sequence *γ*_hHCN2_>*γ*_mHCN1_>*γ*_hHCN3_≍*γ*_mHCN4_.

For mHCN1 channels, the more rapid activation and deactivation compared to mHCN2 allowed us to apply faster pulsing (Fig. 1a) while the remaining analysis was analogue to mHCN2. The illustrated traces at two voltages together with the ensemble average currents show again the characteristic sigmoidal activation time course (top). Notably, the amplitude of the unitary currents was only about half as large as that for mHCN2 channels, resulting in a lower unitary conductance *γ*_mHCN1_=0.84±0.01 pS (Fig. 2a,b; Table S1). Superimposing *cL*_1_(*t*) with *P*_o_(*t*) (Fig. 1a, bottom) reveals that the activation time course is also mainly determined by *cL*_1_(*t*).

The single-channel analysis of the slowly activating mHCN3 and mHCN4 channels required longer hyperpolarizing pulses and intervals at the resting potential (c.f. Extended Data Fig. 1). Hence, both the pulses and the intervals at the resting potential were set to 4 s, resulting in a slow pulsing rate of 0.125 Hz. As an unfavorable consequence, the number of traces per time was much lower and sufficiently many nulls for subtraction were not available. Therefore, single-channel idealization was performed from raw data by a semi-automated procedure (Materials and Methods). The amplitude of the currents were determined as described above.

Unitary mHCN3 currents often started after an exceptionally long first latency and stayed regularly open until the end of the hyperpolarizing pulse (Fig. 1c), yielding the single-channel conductance *γ*_mHCN3_=0.54±0.01 pS. This value is even smaller than that for mHCN1 channels (Fig. 2a,b; Table S1).

A respective analysis was also performed for mHCN4 currents, as demonstrated for a 2-channel patch in Fig. 1d, yielding a single-channel conductance *γ*_mHCN4_=0.51±0.01 pS, a value similar to that in mHCN3 channels (Fig. 2a,b; Table S1). For both, mHCN3 and mHCN4 we also superimposed *cL*_1_(*t*) with P_o_(*t*) (Fig. 1c,d bottom). The activation time courses are mainly determined by *cL*_1_(*t*), indicating that also these channels regularly stay open after they have once opened.

To test whether or not the exceptionally low conductance determined for mHCN4 channels is specific for the mouse, we investigated also the human isoform hHCN4 (Extended Data Fig. 3). All features, the sigmoidal averaged current time course, the matching superimposition of *P_o_*(*t*) with *cL*_1_(*t*) (Extended Data Fig. 3a, bottom) and the single-channel conductance γ_hHCN4_=0.56±0.01 pS (Extended Data Fig. 3b,c; Table S1) resemble those in mHCN3 and mHCN4 channels. The minor difference *γ*_hHCN4_>*γ*_mHCN4_ is viewed to be second-tier and is not further interpreted.

To further substantiate that the single-channel currents reported herein are indeed from expressed mHCN channels, we applied the pertinent channel agonist cAMP (20 μM) to excised patches at single-channel resolution (Extended Data Fig. 4a), a saturating concentration for HCN2 and HCN4 channels ^35,36^. Here, we used patches containing multiple channels with still single-channel resolution to minimize effects of stochastic openings, and we considered the activation time course. For all tested isoforms, mHCN1, mHCN3 and mHCN4, activation was notably accelerated (n=4 for each isoform). The result was verified by three other experiments for each isoform. Among these effects that of mHCN1 is the smallest, confirming previous results ^19^. Because the agonistic effect on mHCN3 conflicts with previous results ^17,20^, we also measured steady-state activation in the absence and presence of 20 μM cAMP. Though the currents were only tiny, we observed a robust shift of activation to positive voltages. Moreover, we confirmed the accelerated activation by fitting an exponential yielding the time constant *τ*_act_ (Extended Data Fig. 5a,b). Together with the previous results on mHCN2 channels ^5,6^, our results demonstrate that cAMP up-regulates all four mHCN1-4 channels leaving the amplitude of the single-channel current unaffected (Extended Data Fig. 4b).

### Molecular mechanism for the different single-channel conductance of HCN channels

We next addressed the question why the conductance of the HCN channels differs up to 3-fold. Alignment of the sequences (Fig. 3a) reveals that for mHCN2, mHCN3, mHCN4 and hHCN4 the two narrow pore regions ^9^, the selectivity filter (SF, CIGYG, green) and the inner gate (I…T…Q, cyan), are identical. This makes it unlikely that the conductance is defined simply by Ohm’s law. We therefore performed mutagenesis in the wider pore region. Its amino-acid identity lies between 83 and 92%. All results are provided by the boxplot in Fig. 3b and Table S2. To improve readability, mHCN1-mHCN4 will be abbreviated in the following by m1-m4.

**Fig. 3.**
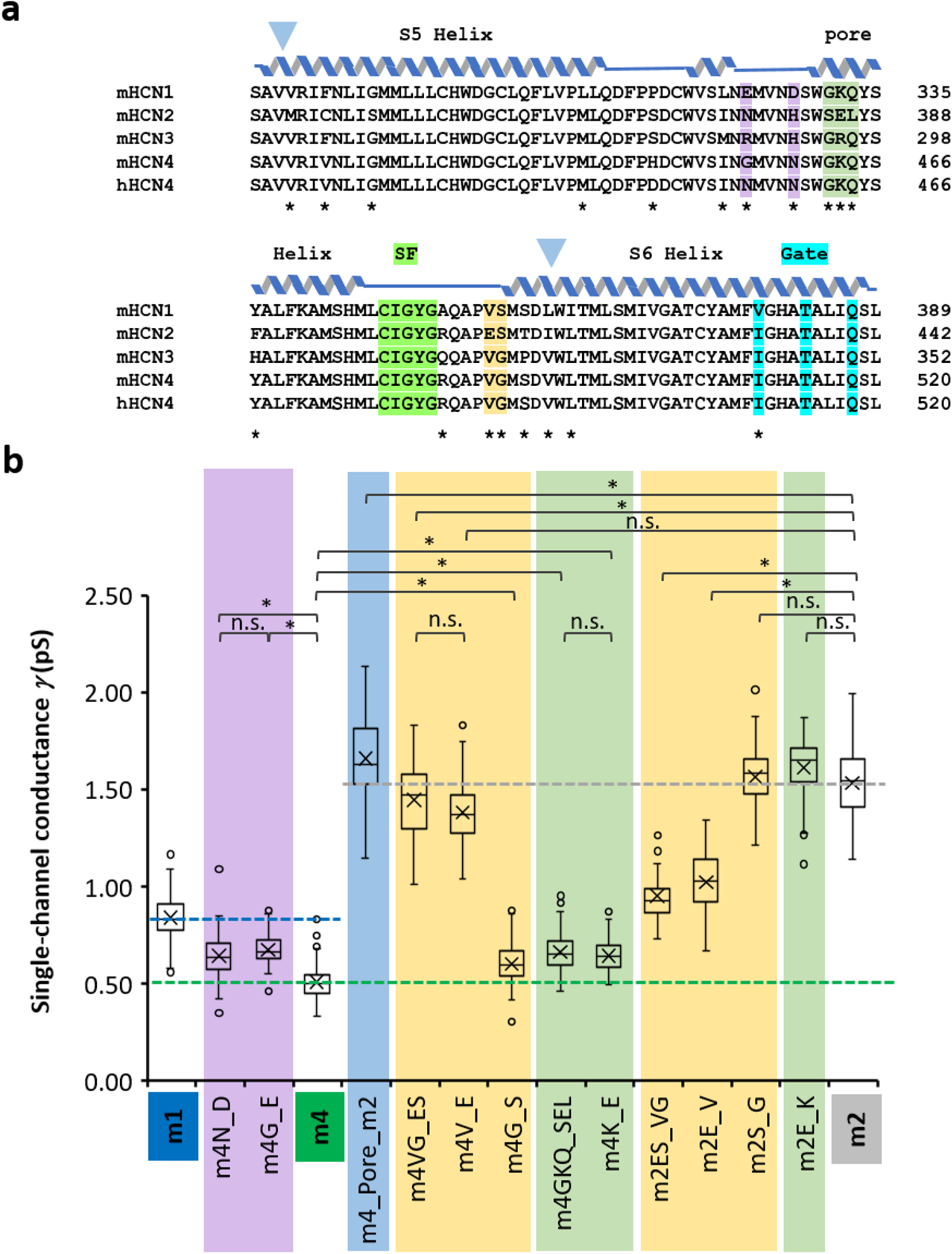
Mutagenesis of key motifs in the pore region. To elucidate why the mHCN channels differ in their conductance by a factor of 3, conspicuous regions in the channel pore were mutated (see also Table S2). **a**, Sequence alignment of the pore regions for the four mouse isoforms and the human hHCN4. Green, selectivity filter (SF); cyan, inner gate. The three motifs differentiating the channels, m4GKQ_m2SEL (brown), m4VG_m2ES (olive), and m4G…N_m1E…D (violet) were mutated. For m4_Pore_m2 the light blue triangles indicate the borders for differences in the pore between m2 and m4. **b**, Boxplot of the unitary conductances of the mutants and relation to the wt channels. The colored dashed lines for the conductances of the wt channels serve as reference. Outliers are included. As in Fig. 2a, the conductance values were calculated from the individual conductance values. Relevant comparisons of the conductance values are indicated by parentheses. * indicates a significant difference with P=0.01; n.s. means not significantly different (pairwise t-tests with Bonferroni-Holm correction). Negative charges in the outer channel vestibule critically modulate the single-channel conductance.

Inserting the whole m2 pore into m4 (m4_Pore_m2) generates the large conductance typical for m2, indicating that the larger conductance is essentially generated by the pore region. The fact that the conductance is even slightly larger than that of m2 suggests an additional small current component by the channel background outside the pore region. It will not be further discussed herein.

We next considered differences between m2 and m4 within the pore region near the SF and the inner gate. Two motifs stand out immediately. In m4, proximal and distal to the SF, a GKQ and VG motif is substituted by a SEL and ES motif in m2, respectively, introducing two negatively charged glutamic acids.

We first transferred the whole ES motif of m2 into m4 and then both amino acids alone (m4VG_ES, m4V_E, m4G_S). The result is that the transfer of a negative glutamate alone generates a large conductance similar to that in m2. Hence, inserting a negative charge in the outer vestibule of the channel (c.f. Fig. 4a,b) essentially augments the conductance. Transferring the second conspicuous m2 motif, SEL, to the respective GKQ position in m4 (m4GKQ_SEL) also augmented the conductance, but to a lesser extent. Transferring the glutamate alone caused an equal effect (m4K_E). To consolidate these effects of the two glutamates, we tested also several reverse m2 mutants where the respective m4 motifs were inserted (m2ES_VG, m2E_V, m2_SG, m2E_K). As expected, m2ES_VG and m2E_V reduced the conductance notably. The fact that the conductance is still some larger than that of m4 suggests that here the still present SEL motif plays a relevant role. As expected in this logic, m2_SG and m2E_K produced a conductance similar to m2. The results so far support the notion that negative charges in the external channel vestibule are responsible for the different conductance values of m2 and m4.

**Fig. 4.**
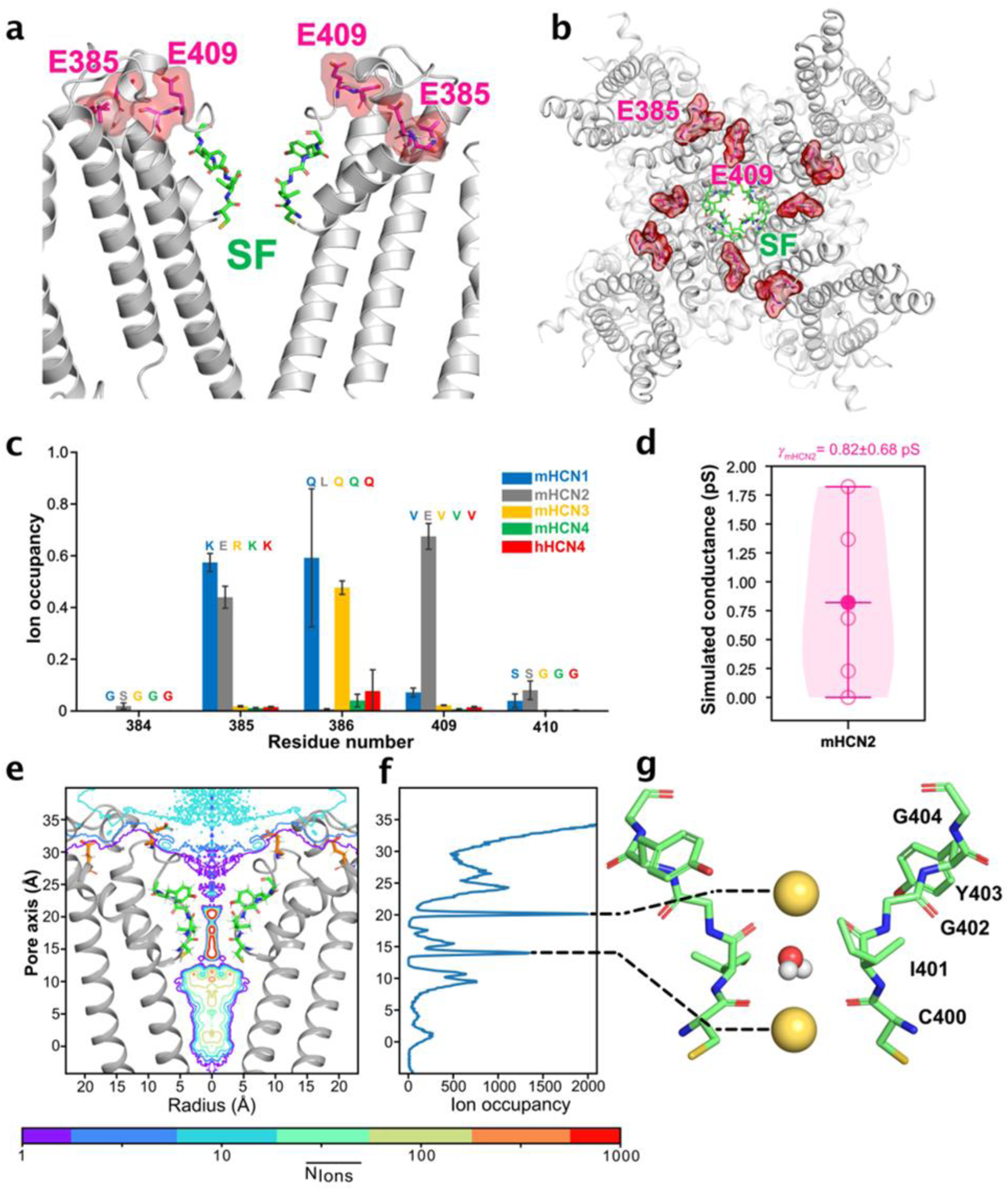
MD simulations of HCN channels. **a**, Side view and **b**, top view of the mHCN2 channel with two key glutamic acid residues highlighted in red and SF residues shown in green. **c**, Residue-wise K^+^ occupancy of the SEL(GKQ,GRK) and ES(VS,VG) motifs derived from the simulations of mHCN1-4 and hHCN4 channels, respectively. Ion occupancy was calculated as the percentage of time the ion is in close proximity (within the first hydration shell of K^+^: 3.4 Å). Three runs of 200 ns without transmembrane potential were performed for each channel. **d**, Conductance derived from simulations of mHCN2, with circles representing the conductance from single simulation runs and the dot representing the average conductance from five parallel simulation runs. Each simulation was performed for 1 µs. **e**, Two-dimensional absolute ion occupancy resolved radially and along the pore axis (z-axis) as a contour plot calculated from five runs of ion permeation simulations when applying a voltage of -700 mV. The orange residues are E385 and E409 as in a. **f**, One-dimensional ion occupancy along the pore axis (z-axis) calculated from five runs of ion permeation simulations at -700 mV. **g**, A snapshot of the mHCN2 SF showing the main populated K^+^ ion configuration and water molecule within the SF.

In the light of this we finally considered the intermediate conductance of m1, a channel carrying the same GKQ and VG motif as m4. The conspicuous difference are two negative charges proximal to the GKQ motif (Fig. 3a). Transferring these charges to the corresponding positions in m4 (m4N_D, m4G_E) augmented the conductance in both m4 mutants in the direction to m1.

The conclusion of these mutagenesis experiments is that the absence or presence of four negative charges in the channel vestibule modulates the single-channel conductance of HCN channels, among which the glutamate in the ES motif of mHCN2 has the strongest effect.

### Differences in ion occupancy of mHCN1-4 in the outer channel vestibule revealed by MD simulations

To mechanistically understand the experimental findings, we conducted atomistic MD simulations on mHCN1-4 and hHCN4 channels. Homology structures of mHCN1-4 and hHCN4 were generated using the open conformation of the rabbit HCN4 channel (PDB ID: 7NMN) ^27^ as template. In this section the numbering of mHCN2 channels is generally used. As shown in Fig. 4a/b, two key glutamic acid residues in the SEL and ES motif identified as responsible for the larger single-channel conductance in the mHCN2 isoform, are located at the outer channel vestibule, with E409 in the ES motif of mHCN2 being closer to the ion conduction pathway.

We first performed MD simulations of mHCN1-4 channels without transmembrane voltage. These simulations revealed distinct residue-wise K^+^ occupancy differences in the SEL(GKQ,GRQ) and ES(VS,VG) motifs between HCN1-4 (Fig. 4c), while the ion occupancy in the rest of the channels remained highly similar. Notably, E409 in mHCN2 showed a very high residue-wise K^+^ occupancy, which was absent at this position in all other HCN isoforms. This finding aligns well with the experimental single-channel data, showing that mHCN2 exhibits the largest single-channel conductance among all HCN isoforms and that introducing a negatively charged glutamic acid residue in the ES motif into mHCN4 substantially increased its conductance.

In addition, we compared the ion occupancy in the SEL(GKQ,GRQ) motif among different HCN isoforms. Here, the mHCN2 channel showed high ion occupancy at residue E385, while HCN1 exhibited high K^+^ occupancy at the corresponding glycine and the neighboring lysine residue (Fig. 4c). We questioned why glycine and lysine could exhibit large cation occupancy despite being neutral and even positively charged, respectively. Analysis of the MD snapshots revealed that K^+^ has a transient ion binding site close to these two residues, enabled by two negatively charged residues, E324 and D328 (Extended Data Fig. 6), that are spatially close to the K332 and adjacent to the SF. These results are again in excellent agreement with the recorded single-channel data, showing that introducing these two negatively charged residues into mHCN4 increased its conductance to match that of mHCN1. In conclusion, the comparison of single-channel data of mHCN1-mHCN4 with atomistic MD simulations confirmed the essential role of ion occupancy differences at the outer channel vestibule in explaining the single-channel conductance differences among the four HCN isoforms.

Since mHCN2 experimentally exhibits the largest single-channel conductance among the four isoforms, we simulated ion permeation through the mHCN2 channel by imposing negative transmembrane voltages during the simulations (Movie 1; details in Materials and Methods). The simulated inward conductance of mHCN2 is 0.82±0.68 pS (Fig. 4d), slightly lower than the experimental single-channel conductance of *γ*_mHCN2_=1.54±0.02 pS. Nonetheless, the simulated conductance of mHCN2 suggested a very low conductance value for the HCN channels, consistent with previously simulated low conductance values of rabbit HCN4 ^37^ and human HCN1 channels ^10,38^. Analysis of one- and two-dimensional ion occupancy of mHCN2 channel along the ion permeation indicated substantial K^+^ occupancy in the outer channel vestibule, partly contributed by the negatively charged E409 pointing toward the ion permeation pathway (Fig. 4e,f). Furthermore, we observed distinct K^+^ binding sites in the SF that are separated by a water molecule (Fig. 4g, Extended Data Fig. 7), similar to the SF ion occupation profiles of a previously simulated rabbit HCN4 channel ^37^ but in sharp contrast to the K^+^ permeation profile in the SF of typical K^+^ channels ^39,40^.

### mHCN2 subunits form heteromeric channels with all three other isoforms

The specific conductance of the mHCN subunits provides a unique opportunity to study heteromerization at the single-channel level because a heteromer with different amounts of negative charges in the vestibule should generate different local concentrations of ions moving to the SF. In principle, when co-expressing two types of subunits (A and B) capable to heteromerize, they can form 2 homotetramers (AAAA, BBBB) and 3 heterotetramers (AAAB, AABB (cis and trans together), ABBB). This would mean that three additional conductance values might appear between those of the homomers. To maximize resolution, we here tested the heteromerization capability of mHCN2 subunits with the other subunits because of its outstanding large conductance. Indeed, sublevels between the homoteramer conductances were observed for all investigated combinations (Fig. 5).

**Fig. 5.**
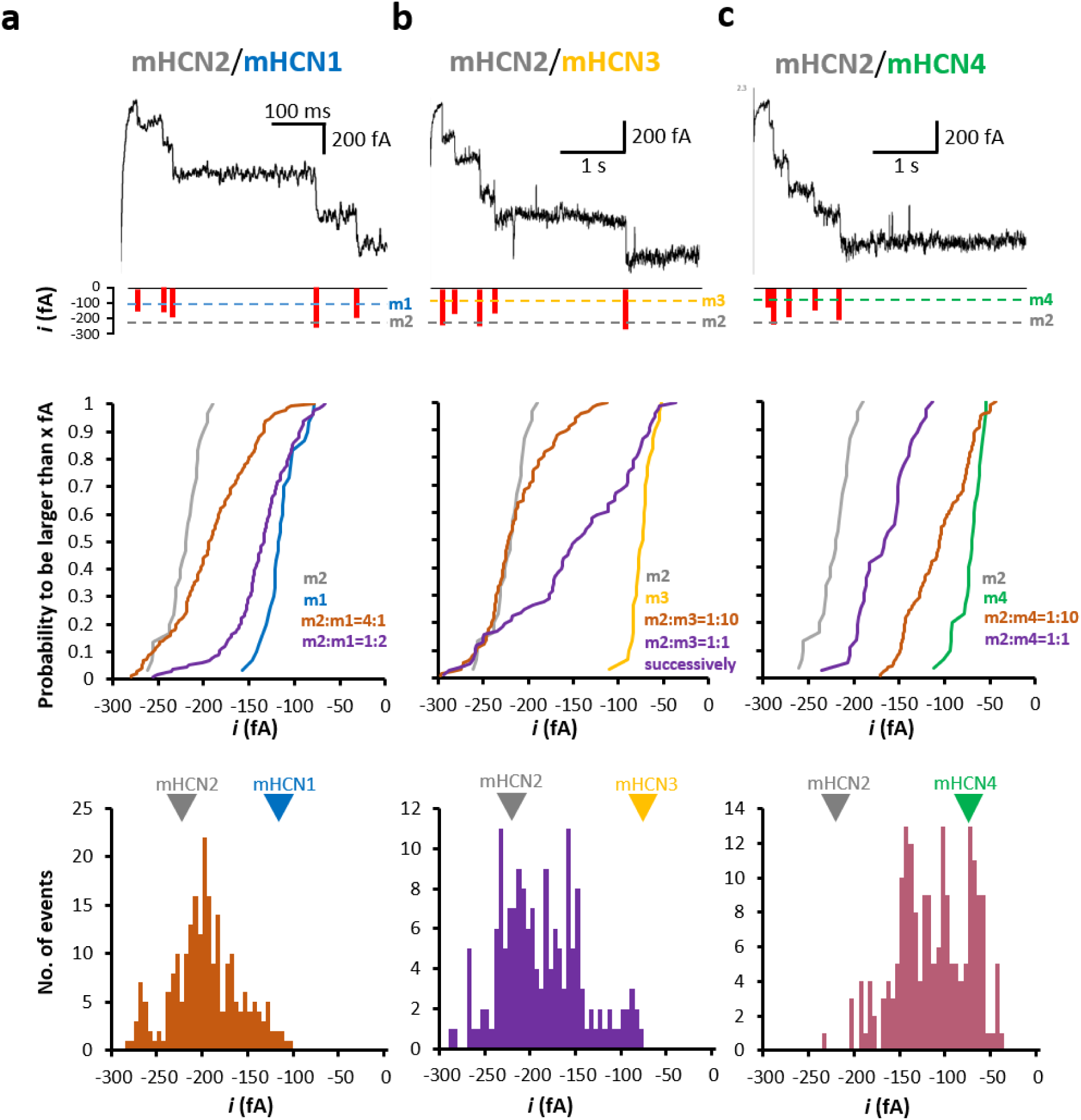
Co-expression of mHCN2 subunits with all other HCN subunits. Different RNA mixtures were generated and injected. The mixtures are specified by ratios of the RNA mass. The voltage was generally -130 mV. On the top a representative trace of a multi-channel patch is shown together with the amplitudes of the individual openings (red bars) determined by amplitude histograms. In the middle, cumulative plots are illustrated for the probability of a single-channel current to be larger than *x* fA, *P*_i>xfA_. At the bottom, the incidence of current levels is plotted as function of the unitary current amplitude. Statistics are provided by Table S3. **a**, Co-expression of m2 and m1. At an RNA ratio m2:m1=1:2, most of the openings were dominated by m1 and while a minority of openings caused either an intermediate or the m2 current level. At an RNA ratio m2:m1=4:1, more intermediate and m2 levels became prominent. The bottom histogram includes 232 events from 60 traces with 6 channels of a single patch to avoid effects of patch variability. **b**, Co-expression of m2 and m3. At m2:m3=1:10 still leaves m2 as dominating subunit. Subsequent RNA injection, m3 at day 0 and m2 at day 6 to 8 (ratio 1:1), resulted in a significant portion of levels between both wt channels. The bottom histogram includes 162 events from 59 traces of a single patch with ∼12 channels to avoid effects of patch variability. **c**, Co-expression of m2 and m4. At m2:m4=1:1 the relationship approximates m2 whereas at m2:m4=1:10 it is closer to m4. The bottom histogram was formed from all 197 events included in the cumulative plot.

These experiments were performed at the voltage of -130 mV only (Fig. 5) and only patches were further analyzed that showed significant levels between those of the respective two wt channels. To best visualize the amplitudes of the single-channel currents of the heteromers between those of the homomers statistically, we preferred cumulative plots of the probability of a single-channel current to be larger than *x* fA, *P*_i>xfA_ (Fig. 5a-c, middle) (For statistics see Table S3). For the wt channels, these *P*_i>xfA_ plots generate steep relationships with clear separation apart from m3 and m4 (Extended Data Fig. 8). The slopes represent variability and resolution of our amplitude measurements.

Co-expressing m2 and m1 in a 1:2 RNA ratio showed that most of the openings were dominated by m1 while a minority generated either intermediate or m2 current levels (Fig. 5a). When increasing the RNA ratio to m2:m1=4:1, clearly more intermediate levels appeared and also m2 levels were more frequent. Co-expression of m2 and m3 in a 1:10 RNA ratio generated a *P*_i>xfA_ relationship dominated by m2 though 10 times more m3 RNA was injected (Fig. 5b). We therefore injected first m3 RNA (day 0) and later (day 6) m2 RNA in a ratio 1:1. Current recordings, performed at day 7, yielded a significant portion of levels between both wt channels, demonstrating that also m3 subunits can heteromerize with m2 subunits. Co-expressing m2 and m4 in a 1:1 RNA ratio generated a *P*_i>xfA_ relationship that was dominated by m2 but also containing a component between the two wt channels (Fig. 5c). When changing the RNA ratio to m2:m4=1:10, the *P*_i>xfA_ relationship shifts to smaller current levels and significantly superimposes with m4, suggesting that the portion of m4 subunits in the heteromers has increased.

To test if the intermediate unitary currents between those of the wt channels group to distinguishable levels (peaks), eventually reflecting different stoichiometries, we built respective histograms (Fig. 5, bottom). The histograms for m2/m4 and m2/m3 reveal indeed discrete peaks, suggesting that an intermediate peak at larger current amplitude is generated by a heteromer with a higher number of m2 subunits due to more external charges in the m2 vestibule. One should be aware, however, that the amplitude of the peaks cannot be interpreted because the incidence of a current level depends also on the expression level of the subunits in a patch and the chance which channel constellation is included. For m2/m1, where the resolution is *a priori* inferior because of the smaller current difference between m2 and m1, distinct levels could not be identified unequivocally (Fig. 5a, bottom).

## Discussion

We showed that all four cloned mouse HCN channel isoforms, functionally identified by the characteristic slow and sigmoidal activation time course, generate a single-channel conductance in the order of 1 pS. Moreover, this single-channel conductance differs up to 3-fold in the sequence *γ*_mHCN2_ > *γ*_mHCN1_ > *γ*_mHCN3_, *γ*_mHCN4_, *γ*_hHCN4_. These tiny conductance values, are in good agreement with results on previous single-channel recordings in both native channels in cardiac sinus node cells ^12,29^ and heterologuously expressed mHCN2 channels ^5,41^. They also confirm values obtained by nonstationary noise analysis in pyramidal neurons ^31^. The simulated conductance of mHCN2 by atomistic MD simulations on the microsecond time scale is also in the order of 1 pS, aligning with the experimental single-channel conductance.

Because the amino acid sequence in the two narrow parts of the pore, the selectivity filter (CIGYG) and the inner gate (I…T…Q), is identical (Fig. 3a) ^9^, the conductance difference must arise from regions outside these narrow pore parts. We identified possible key players and verified them by mutagenesis (Fig. 3b; Table S2). The key players for the larger conductance in the mHCN2 channel compared to the other isoforms are two negatively charged glutamic acids in the outer channel vestibule. In the sequence, one of these negatively charged residues is positioned proximal (SEL motif) and the other distal (ES motif) to the SF (Fig. 4a,b). Notably, we observed excellent agreement between the ion occupancy difference in these two motifs identified by MD simulations and the single-channel conductance, further confirming the essential role of the cation occupancy in the outer vestibule in modulating single-channel conductance of HCN channels. In mHCN1, generating an about 60% larger conductance than mHCN4, two further negative charged were identified, also proximal to the SF, enhancing the ion occupancy at the GKQ motif (corresponding to the SEL motif in mHCN2).

The functional role of rings of negative charges in the channel vestibule to increase the single-channel conductance was extensively studied previously in structurally related voltage-gated sodium channels, generating an inward current of monovalent Na^+^ ions ^42–44^. A proposed role of these charges is to concentrate Na^+^ ions in the vestibule without binding them, supporting a knock-on mechanism for ion permeation ^44^. A similar effect of charges in the channel vestibule on K^+^ ions is likely for HCN channels.

Our co-expression experiments of mHCN2 with either of the other subunits could clearly induce conductance values in-between those of the homomeric channels using different RNA ratios to approximately cover the range between the respective wt channels (Fig. 5, middle). The extreme case was m2/m3. Even in a 1:10 ratio the unitary currents were by far dominated by m2, which again fits to the exceptionally long incubation times of 7-13 days required for m3 alone. To prove the principal capability of the m3 subunits to form heteromers with m2, we used successive RNA injection in a 1:1 ratio (day 0 m3, day 6-8 m2; measurement within 24 hours after m2 injection) and obtained intermediate unitary currents. Though it remains to be shown if such a scenario can appear in nature, it tells that the two types of subunits can assemble. Mechanistically, it is assumed that m3 subunits are produced slowly and linger in the endoplasmic reticulum as monomers or incomplete channels, eventually in complexes with chaperones ^45^. After injection of the m2 RNA, the robust expression of m2 subunits leads to an assembly with the already present m3 subunits.

The pronounced peaks of the intermediate current levels (Fig. 5, bottom) might reflect at least for m2/m4 and m2/m3 channels different stoichiometries (or geometries) of the subunits. This supports the notion that the conductance generated by the m2 subunits with the identified two negative charges in the SEL and ES motif gradually contributes to the overall-conductance depending on the actual number of subunits included. Nevertheless, from our present data it is too early to unequivocally assign the current peaks to the subunit composition. This question can be addressed in future experiments by constructing concatameric HCN channels ^46,47^ with different subunits specifying a defined stoichiometry. Knowing these values it will become possible to analyze single native channels in neurons or SA node cells to learn to what extent these heteromers contribute to the pacemaker current in the physiological context.

### Conclusions

Our analysis of single-channel currents in all four mammalian HCN channels demonstrates that the single-channel conductance is in the range of 1 pS, thereby, however, differing by a factor of up to 3 with *γ*_mHCN2_ > *γ*_mHCN1_ > *γ*_mHCN3_, *γ*_mHCN4_, *γ*_hHCN4_. These differences in the conductance are not generated by the two narrow pore parts, the selectivity filter and the inner gate, but by four negative charges in the outer channel vestibule, leading there to an enrichment of the permeating K^+^ ions. Co-expression of mHCN1, mHCN3 or mHCN4 with mHCN2 demonstrates conductance values in-between those of the respective homomeric channels. Our approach provides essentially new insights into the function of homomeric and heteromeric HCN channels which has great potential to aid developmend of highly specific drugs for the different isoforms and their heteromers.

## Materials and Methods

For standard techniques as oocyte preparation and cRNA injection, molecular biology, two-electrode voltage clamp and patch-clamp recording of macroscopic mHCN3 currents see Supplementary Materials and Methods.

### Patch-clamp recording at single-channel resolution

Currents with single-channel resolution were recorded with the patch-clamp technique from either cell-attached or inside-out patches of *Xenopus* oocytes expressing the respective channels. The patch pipettes were pulled (P-2000 puller, Sutter Instruments, Novato (CA), USA) from thick-walled quartz tubing (outer and inner diameter 1.0 and 0.5 mm, respectively) (Science Products GmbH, Hofheim, Germany) to keep the RC and dielectric noise as low as possible ^48^. The resistance of the pipettes was 7 to 20 MΩ. The bath and pipette solution contained (in mM) 100 KCl, 10 EGTA, 1 MgCl_2_, 10 Hepes (pH 7.2) and 120 KCl, 10 Hepes, 1 MgCl_2_, 1 CaCl_2_ (pH 7.2) respectively. Mg^2+^ was added to elevate patch stability.

In part of the experiments, cAMP (Sigma-Aldrich Corp., St. Louis, USA) was applied to the bath solution to reach, after gently stirring, the saturating concentration of 20 μM.

Particular care was taken to shield the measurements from external interfering: We followed the strategy to position an additional small Faraday box, surrounding the pipette and the bath, in the large Faraday box surrounding the microscope with the headstage (cage-in-cage strategy). With this shielding we could reliably remove all 50 Hz components. Occasional spikes, presumably by inductive sources, could not be completely removed but were sufficiently rare to exclude these traces if required.

Currents were recorded at room temperature with an Axopatch 200B amplifier (Axon Instruments Inc., Foster City (CA), USA). To minimize in inside-out patches effects of the rundown phenomenon typical for HCN channels ^4,12,49^, the measurements were started only 180 seconds after the patch excision. Stimulation and data recording were performed with the ISO3 hard- and software (MFK, Niedernhausen, Germany). The sampling rate was 20 kHz for the faster mHCN1 channels and 5 kHz for the other channels. The on-line filter of the amplifier (4-pole Bessel) was set to 1 kHz. The holding potential was generally -30 mV. The used voltages of the test pulses are indicated in the figures and the diagrams.

The holding potential was generally -30 mV. Hyperpolarizing pulses were sufficiently long to reach significant activation at the end of the pulses. For mHCN1 the pulse duration was 200 ms, for mHCN2 500 ms and for mHCN3, mHCN4 as well as hHCN4 4000 ms. The respective intervals between the pulses were 200 ms, 1500 ms and 4000 ms, respectively. It should be noted that for mHCN1 and mHCN2 the shorter pulses and higher repetition rates allowed us to get much larger numbers of traces than for the slower other channels. This provided the advantage to get empty traces (nulls) which could be subtracted from the individual traces with channel openings to perfectly abolish the capacitive artifacts. Reasonable patches had a seal resistance of >100 GΩ, in many cases 200 to 700 GΩ, or even higher. Such high values were a prerequisite to obtain sufficiently stable conditions for our analyses. These high resistances were not obtained with patches on HEK293 cells.

### Data Analysis

The data of mHCN1 and mHCN2 were generally digital-filtered down to 200 Hz by the filter in the ISO3 software whereas those of mHCN3, mHCN4 and hHCN4 with smaller unitary currents were generally filtered down to 100 Hz. Filtering to the lower frequency of 50 Hz was only performed for the analysis of unitary tail currents at -30 mV for HCN2 channels. The criterion for a channel opening was generally passing the 50% amplitude level.

For mHCN1 and mHCN2 channels, the short pulses provided sufficiently many empty traces without channel activity. These traces were averaged and subtracted from the traces with channel activity, providing the traces in Fig. 1a,b, thereby taking advantage of the ISO3 analysis software to use a sliding null, i.e. five null traces as close as possible to the actual traces were formed and used. This resulted in an improved handling of subtle leak changes during the recording.

In case of the slower channels (mHCN3, mHCN4, hHCN4) the required usage of longer test pulses did not allow us to get as many traces as for the faster channels. We therefore determined the open probability from idealized traces.

To idealize raw traces, the difference between sliding averages preceding and following each current time point were calculated. Points within the expected rise-time, estimated as (3\**f*_c_)^-1^, were omitted. The sliding average window was set to *f*_c_^-1^. Peaks in this difference trace were filtered for artefacts by enforcing minimum (typically 40 fA) and maximum amplitudes (typically 100-300 fA). All traces and gating events were screened and confirmed manually. Note that this strategy has a reduced sensitivity to detect events shorter than *f*_c_^-1^. The strategy was implemented in IgorPRO 7.0.8. (Wavemetrics, Lake Oswege (OR), USA).

The channel number in a patch, *N*, was obtained from the highest number of unitary steps within a trace at -130 mV. The strategy was implemented in IgorPRO 7.0.8. (Wavemetrics, Lake Oswege (OR), USA). The amplitude of the unitary currents was determined locally by individual openings and closures. Because the open and closed times of the channels were reasonably long, it was easy to identify sufficiently many switch transitions from closed to open and open to closed, including also events for the opening of a second and a third channel if present. Such a switch transition contained at least 50 ms before and 50 ms after the switch event. For the analysis of patches with multiple co-expressed channels, also shorter time intervals before and after an opening transition of 15 ms were used. From these trace segments an amplitude histogram was built and fitted by the sum of two, or sometimes three, Gaussian functions (Fig. 1c,d). The chosen bin widths depended on the type of channel and were 12 to 20 fA. In case of the analysis of the unitary tail currents at -30 mV, filtered down to 50 Hz, the bin width was set exceptionally to 6 fA. The fits of the amplitude histograms were performed mainly with the implemented fit routine of the ISO3 software and exceptionally also with the Origin Pro software (2019). The single channel conductance γ of an individual opening was calculated by

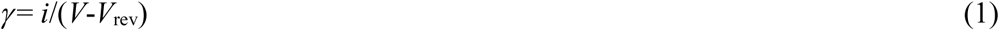

where *i* is the unitary current obtained from a local amplitude histogram, *V* the actual voltage and *V*_rev_ the reversal potential set to 4.6 mV, according to the Nernst equation to

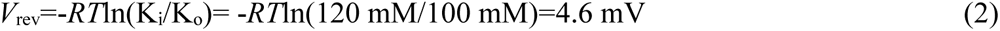

with K_i_ and K_o_ being the K^+^ concentrations in the bath and pipette solution. In cell-attached patches, the resting potential of the oocytes was assumed to be zeroed by the high K_o_ concentration in the bath.

Experimental data are given as mean ± SEM. Boxplots provide the mean (crosses), median (horizontal line), upper and lower quartiles (boxes) and maximum and minimum values (whiskers). Outliers are included.

In time courses of *P*_o_, 95% confidence intervals (Fig. 1a-d, Extended Data Fig. 3a) were estimated from summed idealized traces ^50^.

To enable inclusion of recordings in which clearly visible slow unitary HCN currents were superimposed by large needle-like spikes, presumably evoked by Ca^2+^ activated chloride channels ^51^, we designed a selective median filter to eliminate these needle-like events. This filter was based on the difference between a median filtered (300 points) dataset and the raw data. Only for data points where this difference exceed a threshold, the raw values were replaced by values of the filtered trace. The replaced region was extended (3-13 points) to accommodate the filter frequencies used. Extended Data Fig. 9 illustrates the power of our median filter for a cell-attached patch containing five mHCN2 channels. In case that the median filter was used for displayed recordings, this is indicated in the Fig. legend. No amplitudes were analyzed on positions where data replacement took place.

### Atomistic MD simulation

The homology structure of different HCN channel variants (mHCN1, mHCN2, mHCN3, mHCN4 and hHCN4) were generated based on the cryo-EM structure of the open conformation of homotetrameric rabbit HCN4 channel (PDB ID: 7NMN, ^27^). For the preparation of the molecular dynamic simulation setup, embedding of the entire HCN channels into a 1-Palmitoyl-2-Oleoyl-sn-glycero-3-Phosphocholine (POPC) lipid membrane was carried out in CHARMM-GUI ^52^, where all endogenous ligands were removed before the simulations. All titratable residues of the protein were protonated according to their standard protonation state at pH 7. The simulations were performed with Amber19sb force field ^53^ using TIP3P water model ^54^. All the simulations were performed with GROMACS software package version 2023.3 ^55^ with 900 mM KCl. The system was firstly equilibrated in 6 steps using default scripts provided by the CHARMM-GUI webserver. A time step of 2 fs was used in the simulations of 1.875 ns for equilibration. We prepared two simulation setups. For mHCN1, mHCN2, mHCN3, mHCN4 and hHCN4 simulations, we conducted three independent runs of production simulations without transmembrane voltage, each 200 ns using an integration time step of 2 fs. Besides, we also conducted five independent simulation runs of mHCN2 with each 1000 ns at 700 mV transmembrane voltage by using external electric field ^56^. All simulations details are summarized in Tables S4 and S5. Short-range electrostatic interactions were calculated with a cutoff of 1.0 nm, whereas the long-range electrostatic interactions were treated by the particle mesh Ewald method ^57^. The cutoff for van der Waals interaction was set to 1.0 nm. The simulations were performed at 300 K with an enhanced Berendsen thermostat (GROMACS V-rescale thermostat, ^58^. The Parrinello-Rahman barostat ^59^ was employed to keep the pressure within the system remaining at 1 bar. All bonds were constrained with the Linear Constraint Solver (LINCS) algorithm ^60^. All trajectories were analyzed with GROMACS toolkits and Python3 using MDAnalysis^61^. To calculate the residue-wise ion occupancy, we considered a residue to be interacting with an ion if the distance between them was less than the hydration radius of K^+^, which is 3.4 Å. Molecular visualizations were made with PyMol and Visual Molecular Dynamics (VMD) ^62^.

## Acknowledgment

We like to thank U. Singer, C. Ranke, S. Bernhardt, P. Hachenburg, M. Büschel and M. Händel for the excellent technical support. This work was supported by the Deutsche Forschungsgemeinschaft though the Research Unit 2518 DynIon, project P02, to K.B. and through the CRC1078 ‘Protonation Dynamics in Protein Function’, project C08, to H.S. The authors gratefully acknowledge the computing time made available to them on the Erlangen National High Performance Computing Center (NHR@FAU) and high-performance computer “Lise” at the NHR Center NHR@ZIB.

## Author contributions

K.B. designed the study, did the single-channel recordings, wrote the manuscript and designed most of the figures. U.E. and D.T. performed measurements of macroscopic currents with the TEVC, J.K. measured the cAMP effect on macroscopic currents, R.S. wrote part of the analysis programs including the median filter, H.S. and H.L. performed the MD simulations, analyzed the data and prepared the respective figures, C.S. performed the molecular biology.

