## Supplementary Information for "Subunit-specific conductance of single HCN pacemaker channels at femtosiemens resolution"

by

Klaus Benndorf, Uta Enke, Debanjan Tewari, Jana Kusch, Haoran Liu, Han Sun, Ralf Schmauder, Christian Sattler

### Contents

**Supplementary Results**

mHCN currents in whole *Xenopus* oocytes

**Supplementary Materials and Methods**

Oocyte preparation and cRNA injection

Molecular biology

Two-electrode voltage clamp

Patch-clamp recording of macroscopic mHCN3 currents

**Supplementary References**

**Extended Data Figures 1-9**

**Supplementary Tables S1-S6**

**Movie 1**

### Supplementary Results

##### mHCN currents in whole *Xenopus* oocytes using the TEVC technique

All four mHCN isoforms generated sufficiently large currents (Extended Data Fig. 1a), which is confirmatory for HCN1, HCN2 and HCN4 channels [1-4] but new for HCN3 channels, though for the latter significantly longer expression times of 7-13 days were required than for the other isoforms. Steady-state activation relationships, determined from instantaneous tail currents at -100 mV, were fitted with the Boltzmann function (equation S1), yielding the half-maximum voltage, $V_h$, and the apparent gating charge, $z\delta$ (Extended Data Fig. 1b; Table S6; see Supplementary Methods). $V_h$ shifts to more hyperpolarizing voltages in the sequence mHCN3>mHCN4>mHCN2>mHCN1, thereby covering a voltage range of nearly 40 mV. The value for mHCN3 approximates that for this isoform when expressed in HEK293 cells [5]. If assuming that the endogenous cytosolic cAMP level in the oocytes is 3 to 5 μM [6] and that this is a saturating concentration for all channels, these results presumably reflect activation at saturating cAMP. Activation kinetics for the four mHCN isoforms were quantified by fitting an exponential function (equation S2), yielding the time constant $\tau_{act}$ and a delay time, $t_0$ (Extended Data Fig. 1c,d; see Supplementary Methods). The slowness of activation followed the same sequence mHCN3>mHCN4>mHCN2>mHCN1 as the shift of steady-state activation.

For specifying useful pulse rates in single-channel recordings, we also considered the speed of deactivation at the used holding potential of -30 mV (Extended Data Fig. 1e) which should be short compared to the time interval between hyperpolarizing pulses. At -30 mV, deactivation for mHCN1, mHCN2, mHCN4 and mHCN3 channels was complete after 100 ms, 200 ms, 300 ms and 2 s, respectively. In the single-channel recordings, the time intervals for deactivation were at least double as long to be on the safe side (see Materials and Methods).

### Supplementary Methods

##### Oocyte preparation and cRNA injection

Oocytes were surgically harvested under anesthesia (0.3% 3-aminobenzoic acid ethyl ester) from adult females of *Xenopus laevis*. The procedures regarding the *Xenopus laevis* frogs were approved by the animal ethics committee of the Friedrich Schiller University Jena (UKJ-18-008 from 09 May 2018). The respective protocols were performed in accordance with the approved guidelines. Extreme efforts were made to reduce the stress and to keep the number of frogs to a minimum.

The oocytes were incubated with collagenase A (3 mg/ml, Roche, Grenzach-Wyhlen, Germany) for 105 min in $Ca^{2+}$-free Barth´s solution which contained (in mM) 82.5 NaCl, 2 KCl, 1 $MgCl_2$, 5 HEPES, pH 7.5. Then oocytes at stage IV and V and were manually isolated. They were injected with about 50 ng of cRNA transcribed from the respective coding sequences in pGEM derivatives. We used WT mHCN1 (NM_010408), mHCN2 (NM_008226), mHCN3 (NM_008227), and mHCN4 (NM_001081192) as well as hHCN4 (NM_005477). After injection with cRNA, the oocytes were cultured at 18°C in Barth solution containing (in mM) 84 NaCl, 1 KCl, 2.4 $NaHCO_3$, 0.82 $MgSO_4$, 0.41 $CaCl_2$, 0.33 $Ca(NO3)_2$, 7.5 TRIS, pH 7.4. To gain reasonable expression for the different HCN isoforms, the incubation times were specific: mHCN1: 1-4 days, mHCN2: 1-4 days, mHCN3: 7-13 days, mHCN4: 3-9 days, hHCN4: 3-6 days.

**Molecular biology**

The mouse HCN 1-4 and human HCN 4 genes and all modified subunits were subcloned behind the T7 promoter of pGEMHEnew. Point mutations and the pore exchange were introduced via the overlapping PCR-strategy followed by fragment subcloning using flanking restriction sites. Rightness of the sequences was confirmed by restriction analysis and sequencing (Microsynth SEQLAB, Göttingen, Germany). cRNAs were prepared using the mMESSAGE mMACHINE T7 Kit (Thermo Fisher Scientific, Dreieich, Germany).

**Two-electrode voltage clamp**

Currents in whole oocytes were recorded with the two-electrode voltage clamp (TEVC) technique (OC725C amplifier, Warner Instrument, Hampden, U.S.A.) at room temperature. Microelectrodes were filled with 3 M KCl. Their resistance was 0.3-1 MΩ. The experiments were conducted in ND96 medium containing (in mM) 96 NaCl, 10 Hepes, 2 KCl 1.8 $CaCl_2$ and 1 $MgCl_2$, pH 7.4. The medium was supplemented with 1 mM $BaCl_2$, 100 µM $LaCl_3$ and 100 µM $GdCl_3$ to minimize impact of endogenous channels [7]. The experiments were controlled by the HEKA Patchmaster software (v2x90.5) and LIH8+8hardware (HEKA Elektronik Dr. Schulze GmbH, Lambrecht, Germany). The holding potential was generally -30 mV. Ionic currents were measured with prepulses between -130 and -40 mV, spaced 10 mV, followed by a test pulse to -100 mV used to determine steady-state activation. Because the activation kinetics of mHCN1 is much faster than that of the other isoforms, the prepulse duration for mHCN1 was set to 1 s whereas for the other channels it was set to 4 s.

Steady-state activation was determined from the amplitude of the instantaneous current at the test pulse of -100 mV, measured as mean current of the time interval 5 to 10 ms ( mHCN1) or 10 to 40 ms (other tested HCN isoforms) after the begin of the test pulse. The relative amplitude of the tail current, $I/I_{max}$, was determined by relating the actual amplitude $I$ of the instantaneous current to $I_{max}$ following the prepulse to -130 mV. This relationship was plotted as function of voltage and fitted by the Boltzmann function

$$I/I_{max}=1/(1+\exp(-z\delta F(V-V_h)/RT)) \qquad \text{(S1)}$$

yielding the midpoint voltage $V_h$ of half-maximum activation and the equivalent gating charge $z\delta$. $R$ is the molar gas constant, $T$ the temperature in Kelvin (K), and $F$ the Faraday constant.

The time course of activation was quantified by a time constant $\tau_{act}$ that was obtained by fitting the sum an exponential function to the time courses which included a delay interval, $t_0$, according to

$$I(t) = \begin{cases} I_{inst} & , for\ (t < t_0) \\ A \times \{1 - \exp[-(t - t_0)/\tau_{act}] + I_{inst}\} & , for\ (t \geq t_0) \end{cases} \qquad \text{(S2)}$$

where $A$ and $I_{inst}$ are the amplitude of the time-dependent and time-independent (leak plus instantaneous current), respectively.

Fits were performed by IgorPRO (Wavemetrics, Lake Oswege (OR), USA) or OriginPro 2016G software (OriginLab Corporation, Northampton, MA, USA).

**Patch-clamp recording of macroscopic mHCN3 currents**

For determining steady-state activation of mHCN3 channels with and without 20 μM cAMP, macroscopic currents were recorded from inside-out patches. The patches were positioned in front of solution outlets for either control bath solution without and with cAMP. The pipette resistance was 1.7 to 2.4 MΩ. The other recording conditions were similar to the single-channel recordings. Steady-state activation was measured analogue to the TEVC recordings by a double pulse protocol in which the hyperpolarising pulses of 4 s duration were followed by a pulse to -100 mV. The normalized amplitude of the instantaneous current at -100 mV was plotted as function of voltage and fitted by equation (S1). Due to the exceptional low expression density, even after more than 7 days incubation time, it seemed that reasonable macroscopic recordings were only obtained from large patches which were presumably located in the pipette interior. This seemed to impede the effectivity of the solution exchange which might have led to an underestimation of the cAMP-induced shift of $V_h$.

### Extended Data Figures

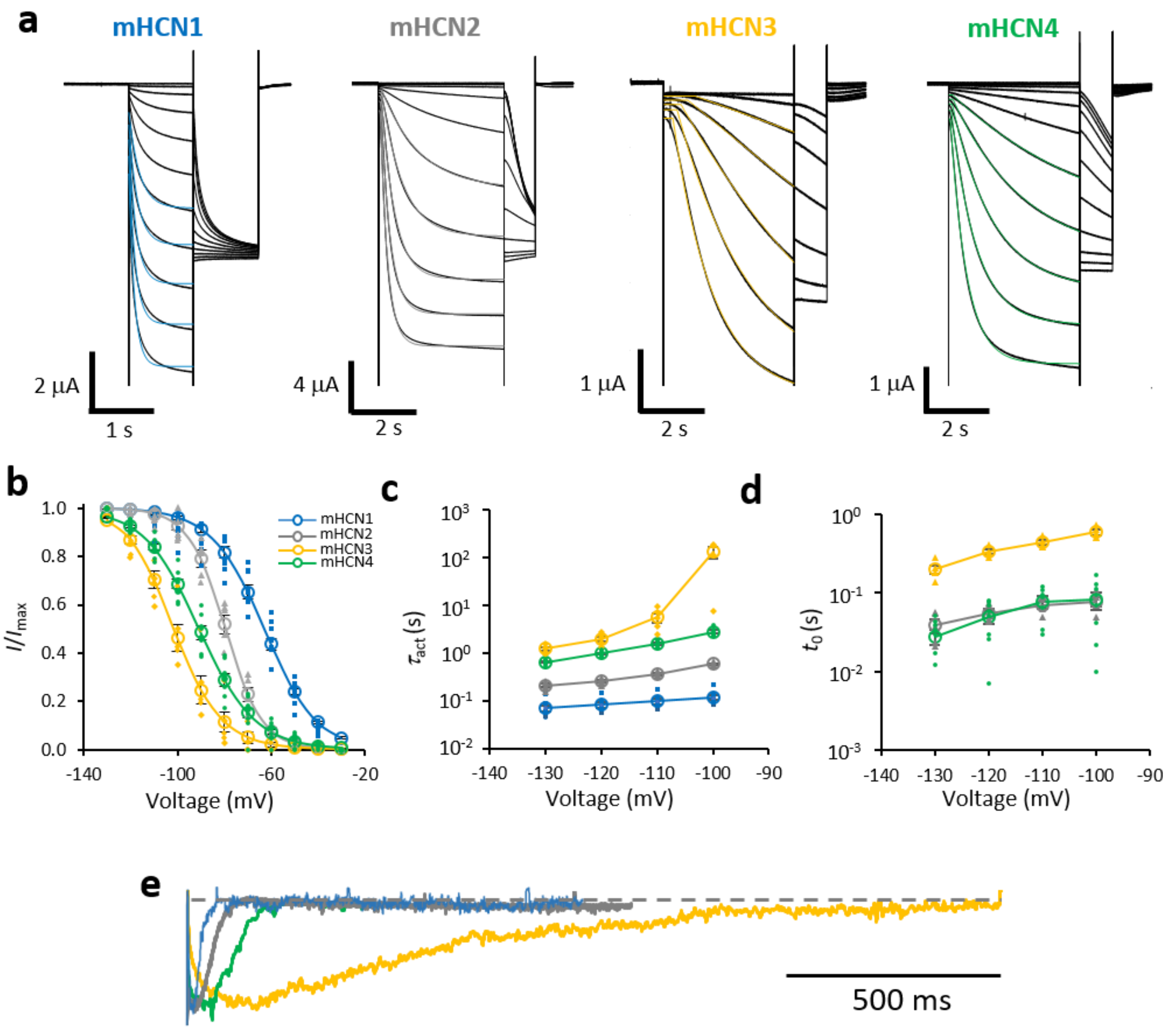

**Extended Data Fig. 1 | Macroscopic currents of the isoforms mHCN1 to mHCN4. a**, Representative currents recorded with the TEVC technique with double pulse protocols. Holding potential -30 mV; potentials first pulse -30 to -130 mV spaced 10 mV; potential second pulse -100 mV. The instantaneous current amplitude at -100 mV was used to construct steady-state activation relationships. **b**, Steady-state activation relationships. Means of 5-8 individual recordings are shown. The errors indicate SEM. The data points were fitted with the Boltzmann function (equation S1) yielding: mHCN1, $V_h$=-63.1 ± 1.1 mV, $z\delta$=2.3 ± 0.1; mHCN2, $V_h$=-79.5 ± 1.3 mV, $z\delta$=3.3 ± 0.3; mHCN3, $V_h$=-99.9 ± 2.0 mV, $z\delta$=2.5 ± 0.1; mHCN4, $V_h$=-90.5 ± 1.3 mV, $z\delta$=2.2 ± 0.1. **c**, Activation time constant, $\tau_{act}$ as function of the voltage. $\tau_{act}$ was obtained from fits of an exponential function with a delay ($t_0$) to the activation time courses (Materials and Methods; equation S2). **d**, Delay, $t_0$, as function of voltage. For mHCN1 the delay was too short for a valid determination. **e**, Tail current kinetics. Representative tail currents of ensemble currents at -30 mV following pulses to -130 mV obtained from macropatches. The current amplitudes were normalized with respect to the peak values to ease comparison of the different kinetics.

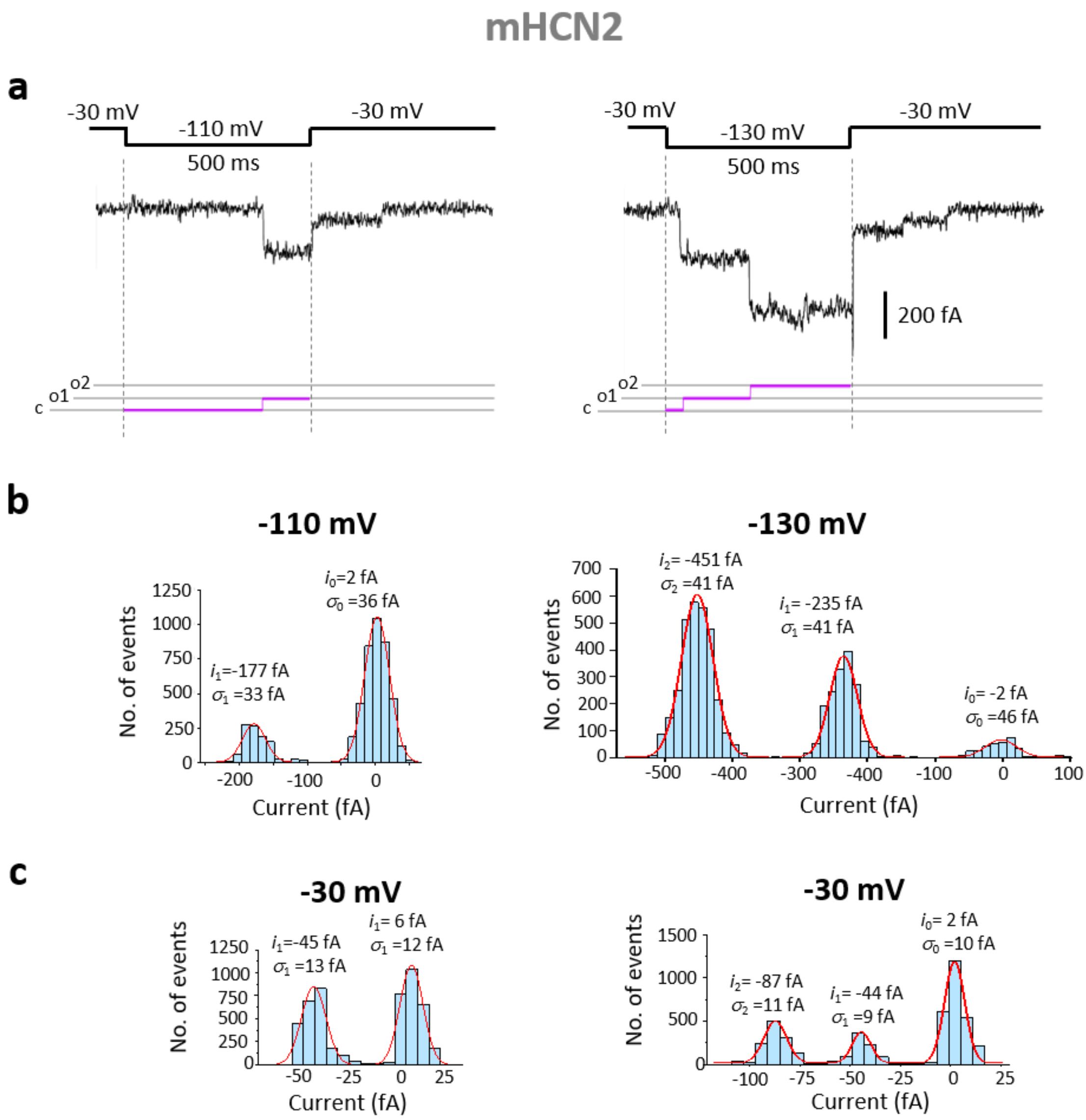

**Extended Data Fig. 2 | Determination of the amplitude of unitary currents. a,** Representative traces of a two-channel patch at -110 and -130 mV. Filter 200 Hz. **b**, Amplitude histograms for the traces in a. If not otherwise noted, the histograms were formed for individual openings >50 ms before and >50 ms after an opening transition. Bin width 12 fA. **c**, Amplitude histograms for the tail currents. Filter 50 Hz. Bin width 6 fA. All histograms were fitted in the usual way with respective sums of Gaussian functions yielding the indicated values for the single-channel current, $i_x$, and the standard deviation, $\sigma_x$, of the levels.

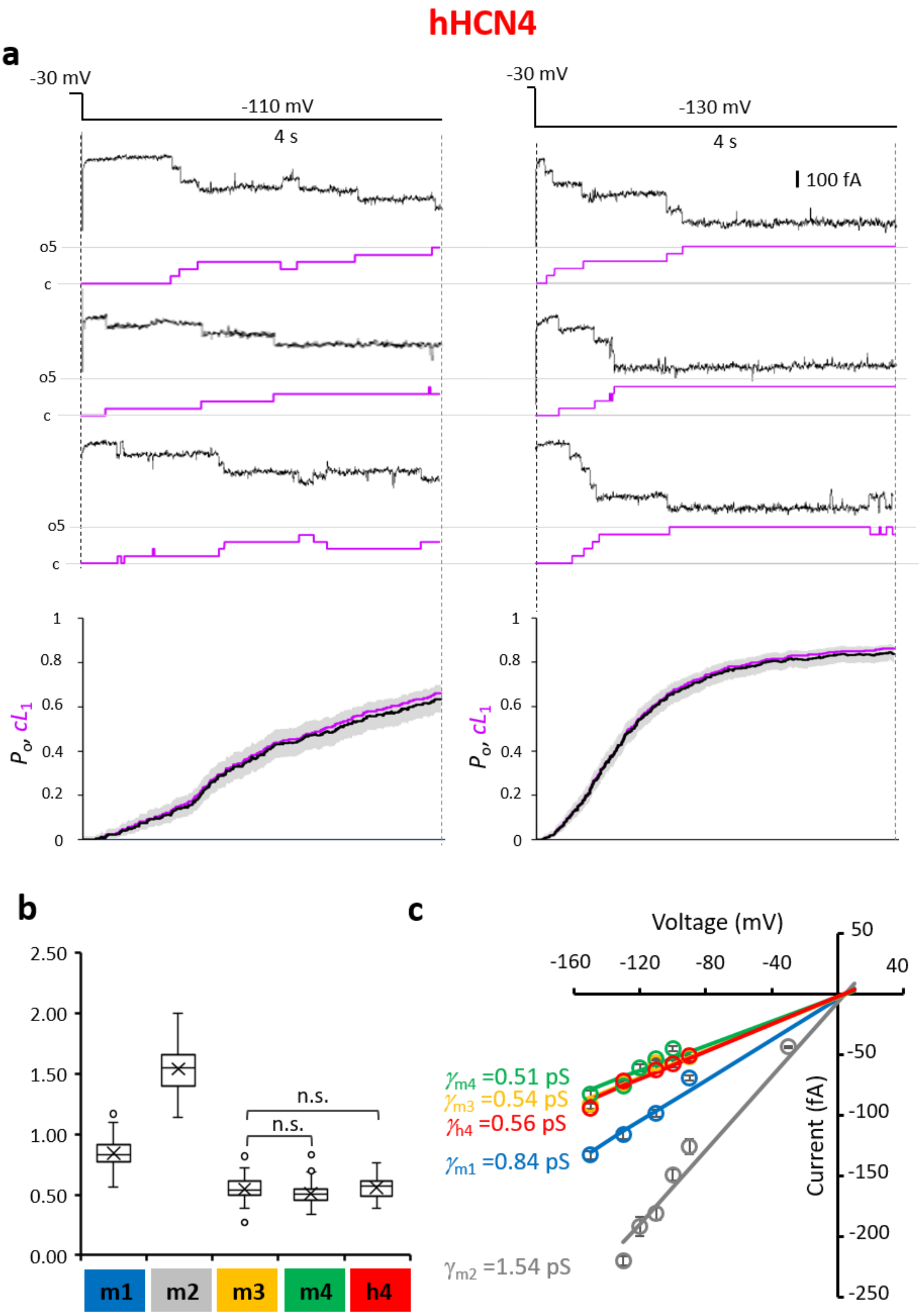

**Extended Data Fig. 3 | Single-channel currents of human hHCN4 channels. a.** Representative recordings of a 5-channel patch at -110 mV and -130 mV. Raw traces. Filter 100 Hz. As for the mouse isoforms, below each trace the corresponding idealized trace is indicated in magenta; c: closed level; o5: open level 5. Superimposition of $cL_1$(t) (magenta) with $P_o$(t) (black) is shown at the bottom as obtained from 46 (-110 mV) and 75 (-130 mV) traces. $cL_1$(t) approximately matches $P_o$(t). **b**, Boxplot of the unitary conductances of mHCN1-mHCN4 (Fig. 2a) including the results for hHCN4. As for the other channels, also the values for hHCN4 were calculated from the individual conductance values. The numbers of included openings, $n_{op}$, and patches, $n_p$, for hHCN4 are also provided by Table S1. **c**, *iV*-relationships for the unitary currents as function of voltage for hHCN4 compared to mHCN1-4 and. hHCN4 channels show a similarly low conductance as mHCN3 and mHCN4 channels.

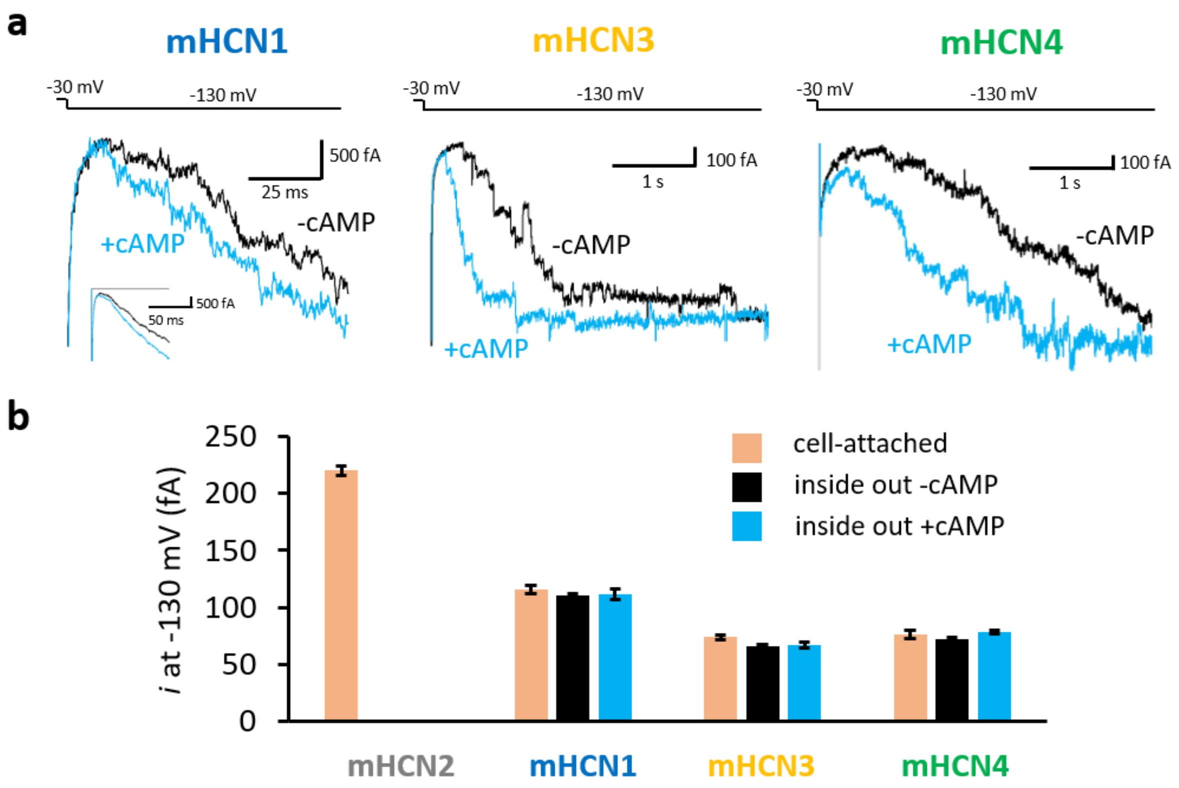

**Extended Data Fig. 4 | cAMP accelerates activation of single mHCN1, mHCN3 and mHCN4 channels.** All measurements were performed in inside-out patches containing multiple channels with single-channel resolution at -130 mV. **a**, Representative traces. Traces in the absence and presence of 20 µM cAMP were recoded from the same inside-out patch. mHCN1: Filter 200 Hz. The inset for mHCN1 channels illustrates ensemble average currents of 15 successive traces of the same patch to illustrate that the cAMP effect is only decent. mHCN3: Patch with ~15 channels. Filter 100 Hz. mHCN4: Patch with ~20 channels, Filter 100 Hz. **b**, Comparison of the single-channel conductance in cell-attached patches, inside-out patches without cAMP and inside-out patches with cAMP. The error bars indicate SEM of 10 to 29 individual openings from 3 to 5 patches. The conductance does not depend on the patch configuration.

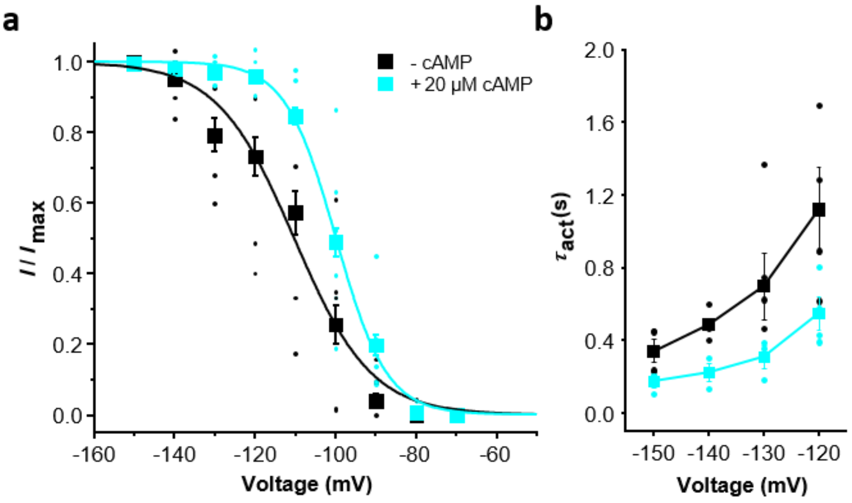

**Extended Data Fig. 5 | Effect of cAMP on the gating of mHCN3 channels in macroscopic currents. a,** Steady-state activation in the absence and presence of 20 μM cAMP. The data points (means and individual points) were fitted by equation (S1) yielding the following parameters: control: $V_h$= -110.1±5.1mV (n=5); +cAMP: $V_h$= -100.2±2.3mV (n=6). **b**, Time constant of activation, $\tau_{act}$, as function of voltage. $\tau_{act}$ was obtained by fitting a single exponential to the activation time course of a current.

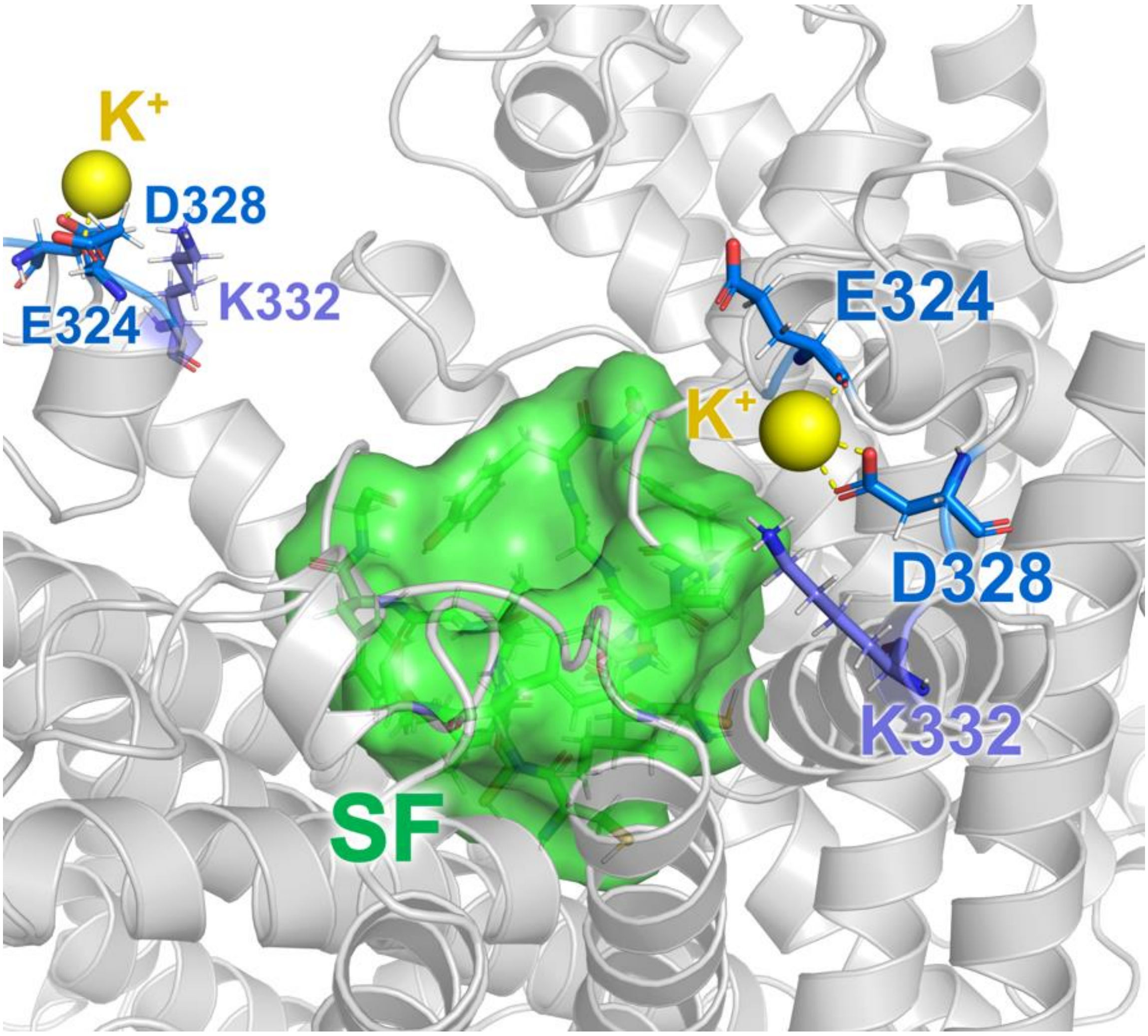

**Extended Data Fig. 6 | Additional $K^+$ binding site at the outer channel vestibule of a mHCN2 channel enabled by the negatively charged E324 and D328 residues.** This binding site was predicted by the MD simulations without transmembrane voltages. Only two subunits are shown for clarity. Green: Selectivity filter.

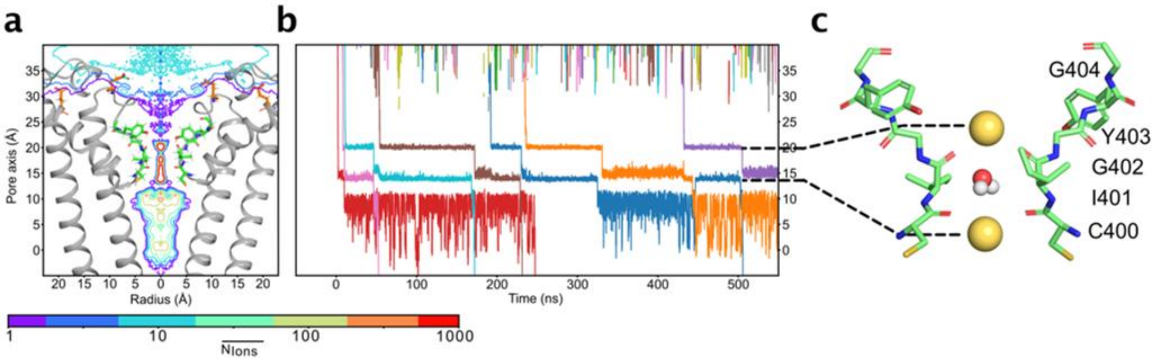

**Extended Data Fig. 7 | $K^+$ occupancy and major binding sites along the ion conduction pathway in a mHCN2 channel. a,** Two-dimensional ion occupancy resolved radially and along the pore axis (z-axis) as a contour plot calculated from five runs of ion permeation simulations. **b**, Representative traces of $K^+$ passing through the SF of the mHCN2 channel pore with each permeating $K^+$ shown in different color. **c**, Snapshot of the mHCN2 SF showing the main populated $K^+$ ion configuration and water molecule within the SF. 5 runs of 1 μs simulations were performed for a mHCN2 channel at 700 mV transmembrane voltage by using an external electric field.

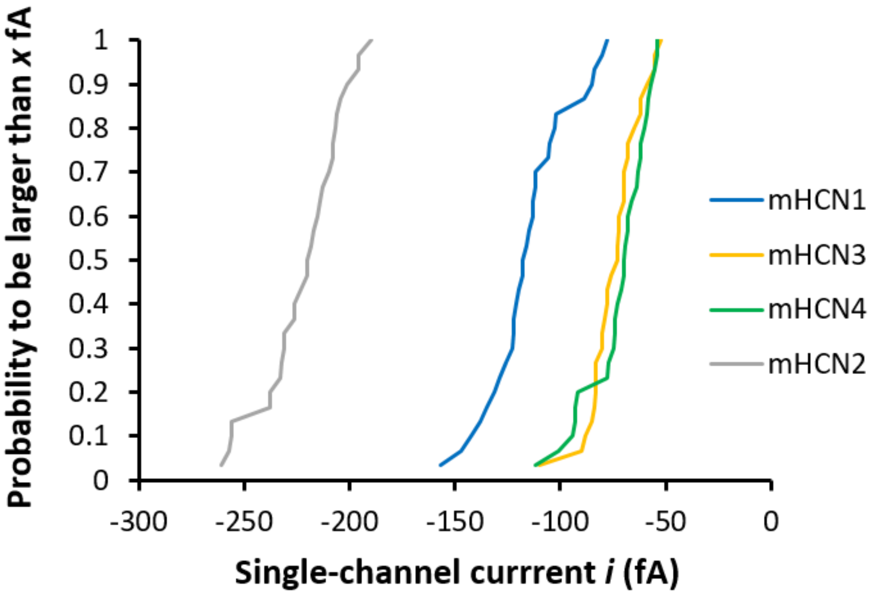

**Extended Data Fig. 8 | Cumulative plot of the single-channel current amplitude of wt channels.**

Plotted is the probability, $P_{i>xfA}$, that a single-channel current is larger than *x* fA. Statistics are provided by Table S3.

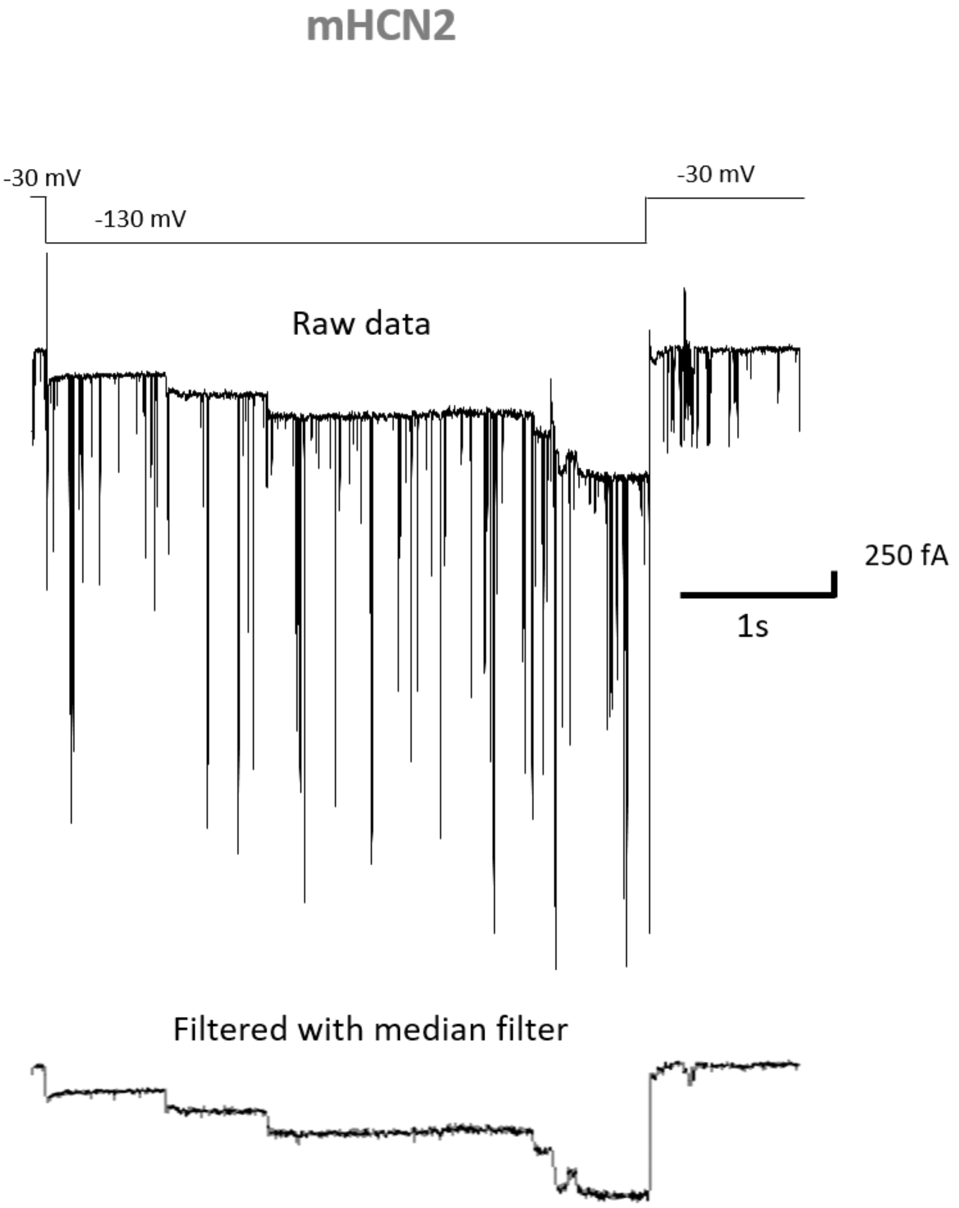

**Extended Data Fig. 9 | Median filter eliminates chloride channels and preserves unitary HCN current events.** Cell-attached patch containing five mHCN2 channels. **a**, Trace with overlapping chloride channels. **b**, Same trace as in a after applying the median filter.

### Supplementary Tables

| Wild-type channels | | | | |
|---|---|---|---|---|
| Isoform | Short name | $\gamma$ (pS) | $n_p$ | $n_{op}$ |
| mHCN1 | m1 | 0.84±0.01 | 6 | 77 |
| mHCN2 | m2 | 1.54±0.02 | 5 | 61 |
| mHCN3 | m3 | 0.54±0.01 | 7 | 62 |
| mHCN4 | m4 | 0.51±0.01 | 5 | 64 |
| hHCN4 | h4 | 0.56±0.01 | 10 | 75 |

**Table S1 | Parameters for single wt channels.** The conductance values were determined as means and SEM of the individual conductance values as specified in Materials and Methods. $n_p$ and $n_{op}$ specify the number of patches and total number of openings included in the analysis, respectively.

| Mutants | | | | | |
|---|---|---|---|---|---|
| Motif | Short name | Systematic name | $\gamma$ (pS) | $n_p$ | $n_{op}$ |
| Pore region | m4_Pore_m2 | mHCN4V416-V492_mHCN2M338-I414 | 1.66±0.02 | 5 | 97 |
| m4VG-m2ES | m4VG_ES | mHCN4V487E_G488S | 1.45±0.02 | 6 | 124 |
| | m4V_E | mHCN4V487E | 1.38±0.02 | 5 | 88 |
| | m4G_S | mHCN4G488S | 0.60±0.01 | 3 | 63 |
| | m2ES_VG | mHCN2E409V_S410G | 0.95±0.02 | 3 | 42 |
| | m2E_V | mHCN2E409V | 1.02±0.02 | 4 | 46 |
| | m2S_G | mHCN2S410G | 1.56±0.02 | 3 | 60 |
| m4GKQ-m2SEL | m4GKQ_m2SEL | mHCN4G462S_K463E_Q464L | 0.67±0.01 | 9 | 109 |
| | m4K_E | mHCN4K463E | 0.65±0.01 | 3 | 72 |
| | m2E_K | mHCN2E385K | 1.61±0.02 | 3 | 77 |
| m1G…N-m4E…D | m4G_E | mHCN4G455E | 0.68±0.01 | 4 | 70 |
| | m4N_D | mHCN4N459D | 0.64±0.01 | 3 | 65 |

**Table S2 | Parameters for mutant channels.** The conductance values were determined as means and SEM of the individual conductance values as specified in Materials and Methods. The abbreviations correspond to Fig. 3. $n_p$ and $n_{op}$ specify the number of patches and total number of openings included in the analysis, respectively.

| | | Short name | Mixture | $n_p$ | $n_{ch}$ | $n_{op}$ |
|---|---|---|---|---|---|---|
| **Wild-type channels** | mHCN1 | m1 | | 6 | >10 | 30 |
| | mHCN2 | m2 | | 5 | >15 | 30 |
| | mHCN3 | m3 | | 6 | >20 | 30 |
| | mHCN4 | m4 | | 4 | >20 | 30 |
| **Co-expressions** | mHCN2 and mHCN1 | m2:m1 | 1:2 | 2 | >19 | 134 |
| | | | 4:1 | 3 | >25 | 150 |
| | mHCN2 and mHCN3 | m2:m3 | 1:10 | 2 | >25 | 123 |
| | | | 1:1 subsequent | 5 | >21 | 76 |
| | mHCN2 and mHCN4 | m2:m4 | 1:1 | 3 | >7 | 36 |
| | | | 1:10 | 2 | >11 | 161 |

**Table S3 | Parameters for cumulative plots of unitary currents.** The data refer to Fig. 5, middle, and Extended Data Fig. 8. $n_p$, $n_{ch}$ and $n_{op}$ indicate for each constellation of co-expression the number of included patches, the total number of channels included in these $n_p$ patches and the number of individual openings, respectively. In multichannel patches, only up to the first 7 overlapping openings were used. An opening was only used if both the time interval before the openings step and after the opening step exceeded 20 ms. Events whose amplitude exceeded 300 fA (maximum amplitude of m2) were discarded by inspection with the assumption that they were generated by more than one channel. The number of channels included in the available patches, $n_{ch}$, is given as minimum because in multi-channel patches its specification becomes increasingly vague in proportion to its number.

| Channels | Cation | Force field | C (mM) | V (mV) | T (K) | Replicates | Simulation time (ns) |
|---|---|---|---|---|---|---|---|
| mHCN1 | $K^+$ | Amber 19SB | 900 | 0 | 300 | 3 | 200 |
| mHCN2 | $K^+$ | Amber 19SB | 900 | 0 | 300 | 3 | 200 |
| mHCN3 | $K^+$ | Amber 19SB | 900 | 0 | 300 | 3 | 200 |
| mHCN4 | $K^+$ | Amber 19SB | 900 | 0 | 300 | 3 | 200 |
| hHCN4 | $K^+$ | Amber 19SB | 900 | 0 | 300 | 3 | 200 |
| mHCN2 | $K^+$ | Amber 19SB | 900 | -700 | 300 | 5 | 1000 |

**Table S4 | List of simulation systems.**

| Channels | Number of protein atoms | Number of water molecules | Number of POPC | Num ber of $K^+$ | Num ber of $Cl^-$ | System atoms in total |
|---|---|---|---|---|---|---|
| mHCN1 | 32172 | 69920 | 437 | 1079 | 1079 | 302648 |
| mHCN2 | 32296 | 69920 | 437 | 1080 | 1088 | 302782 |
| mHCN3 | 32228 | 69600 | 435 | 1073 | 1081 | 301472 |
| mHCN4 | 32380 | 69440 | 434 | 1072 | 1096 | 301024 |
| hHCN4 | 32364 | 69440 | 434 | 1072 | 1084 | 300996 |
| mHCN2 (with voltage) | 32296 | 69920 | 437 | 1080 | 1088 | 302782 |

**Table S5 | Simulation system details.**

| *V* (mV) | mHCN1 (n=8) | | mHCN2 (n=5) | | mHCN3 (n=5) | | mHCN4 (n=8) | |
|---|---|---|---|---|---|---|---|---|
| | $\tau_{act}$ (s) | $t_0$ (s) | $\tau_{act}$ (s) | $t_0$ (s) | $\tau_{act}$ (s) | $t_0$ (s) | $\tau_{act}$ (s) | $t_0$ (s) |
| -90 | 0.147 ± 0.020 | -0.006 ± 0.002 | 1.000 ± 0.054 | 0.077 ± 0.013 | 206.113 ± 12.733 | 0.800 ± 0.056 | 5.894 ± 0.776 | 0.073 ± 0.032 |
| -100 | 0.117 ± 0.016 | -0.008 ± 0.002 | 0.580 ± 0.026 | 0.078 ± 0.010 | 128.412 ± 26.856 | 0.759 ± 0.108 | 2.481 ± 0.094 | 0.081 ± 0.020 |
| -110 | 0.097 ± 0.013 | -0.008 ± 0.002 | 0.364 ± 0.018 | 0.071 ± 0.008 | 6.440 ± 1.348 | 0.495 ± 0.052 | 1.560 ± 0.090 | 0.077 ± 0.013 |
| -120 | 0.081 ± 0.011 | -0.008 ± 0.002 | 0.257 ± 0.015 | 0.055 ± 0.007 | 2.157 ± 0.263 | 0.351 ± 0.027 | 0.961 ± 0.046 | 0.049 ± 0.009 |
| -130 | 0.070 ± 0.011 | -0.007 ± 0.002 | 0.208 ± 0.018 | 0.039 ± 0.007 | 1.289 ± 0.121 | 0.205 ± 0.021 | 0.640 ± 0.030 | 0.028 ± 0.006 |

**Table S6 | Parameters for the analysis of the macroscopic currents recorded with the TEVC technique.** The activation time course were fitted with equation (S2) yielding an initial delay, $t_0$, and a time constant for activation, $\tau_{act}$, both given as mean ± SEM. The negative $t_0$ values indicate that the delay was too short for a reasonable analysis. These values were not further interpreted.

### Movies

**Movie 1. $K^+$ permeation in mHCN2 channel during a 250 ns trajectory.** The simulation was performed using AMBER19sb force field with an external electric field of -700 mV. For clarity, only two subunits of mHCN2 are shown in cartoon (violet), the SF is shown in green sticks, and E385 and E409 in magenta sticks. Permeating $K^+$ ions are shown in spheres of various colors, while the rest of $K^+$ ions in white. Water molecules are drawn as blue O bonded to white H atoms. Z-positions of permeating $K^+$ ions along the pore axis are monitored during ion permeation.
